# Mechanosensing and Sphingolipid-Docking Mediate Lipopeptide-Induced Immunity in *Arabidopsis*

**DOI:** 10.1101/2023.07.04.547613

**Authors:** Jelena Pršić, Guillaume Gilliard, Heba Ibrahim, Anthony Argüelles-Arias, Valeria Rondelli, Jean-Marc Crowet, Manon Genva, W. Patricio Luzuriaga-Loaiza, Estelle Deboever, M. Nail Nasir, Laurence Lins, Marion Mathelie-Guinlet, Farah Boubsi, Sabine Eschrig, Stefanie Ranf, Stephan Dorey, Barbara De Coninck, Thorsten Nürnberger, Sébastien Mongrand, Monica Höfte, Cyril Zipfel, Yves F. Dufrêne, Alexandros Koutsioubas, Paola Brocca, Magali Deleu, Marc Ongena

## Abstract

Bacteria-derived lipopeptides are immunogenic triggers of host defenses in metazoans and plants. Root-associated rhizobacteria produce cyclic lipopeptides that activate systemically induced resistance (IR) against microbial infection in various plants. How these molecules are perceived by plant cells remains elusive. Here, we reveal that immunity activation in *Arabidopsis thaliana* by the lipopeptide elicitor surfactin is mediated by docking into specific sphingolipid-enriched domains and relies on host membrane deformation and subsequent activation of mechanosensitive ion channels. This mechanism leads to host defense potentiation and resistance to the necrotroph *B. cinerea* but is distinct from host pattern recognition receptor-mediated immune activation and reminiscent of damage-induced plant immunity.

## Main Text

Lipopeptides (LPs) represent a prominent and structurally heterogeneous class of molecules among the broad spectrum of small specialized metabolites synthesized by bacteria. Besides serving key functions for the ecological fitness of the producer (motility, biofilm formation, colonization, nutrient acquisition, or antagonism towards competing neighbors), some LPs also act as triggers of immune responses that restrict pathogen infection of metazoans and plants^1, 2^. The vast majority of LPs formed by plant-associated bacteria are comprised of a partly or fully cyclized oligopeptide linked to a single fatty acid chain. Some of these cyclic lipopeptides (CLP) formed by beneficial species belonging to the *Pseudomonas* and *Bacillus* genera are potent elicitors of immune responses in the host plant leading to a systemically induced resistance (IR) against infection by microbial pathogens^2, 3^. This CLP-induced plant resistance is a key process for biocontrol of crop diseases ^4^, but, in contrast to pattern-triggered immunity (PTI), its molecular basis remains poorly understood. Like in animals, PTI in plants relies on the detection of specific molecular motifs (Microbe-Associated Molecular Patterns (MAMPs) via cell-surface plasma membrane (PM)-localized Pattern-Recognition Receptors (PRRs)^5^. Upon assembly of higher order receptor complexes involving conserved co-receptors, PRRs activate receptor-like cytoplasmic kinases (RLCKs) such as BIK1 and its closest homolog PBL1 described as key convergent signaling hubs. This leads to phosphorylation of numerous substrate proteins and subsequent induction of a well-characterized immune response^6^. Early hallmarks of PTI signaling in plants include apoplastic burst of reactive oxygen species ([ROS]_apo_), calcium influx, medium alkalinization indicating H^+^/K^+^ exchange and membrane depolarization, MAPK phosphorylation cascade and initiation of transcriptional reprogramming ^7–10^.

The CLP surfactin (Srf, **Fig 1A**) is well conserved in plant beneficial bacilli^11^ and is among the bacterial compounds best described as immunity elicitor in several plant species^2^. In *Arabidopsis thaliana* ecotype Col-0 (hereafter, *Arabidopsis*), root treatment with purified Srf (at 10 µM as minimal active concentration previously determined^12^ and used as a mix of naturally produced homologues slightly differing in the length of the fatty acid tail, see **Suppl Fig 1**) triggers IR and significantly reduces leaf infection by the grey mold pathogen *Botrytis cinerea* (**Fig 1B**). Therefore, we used Srf as a model to further investigate the molecular mechanisms determining CLP perception and immunity stimulation in *Arabidopsis* root cells.

**Fig. 1.**
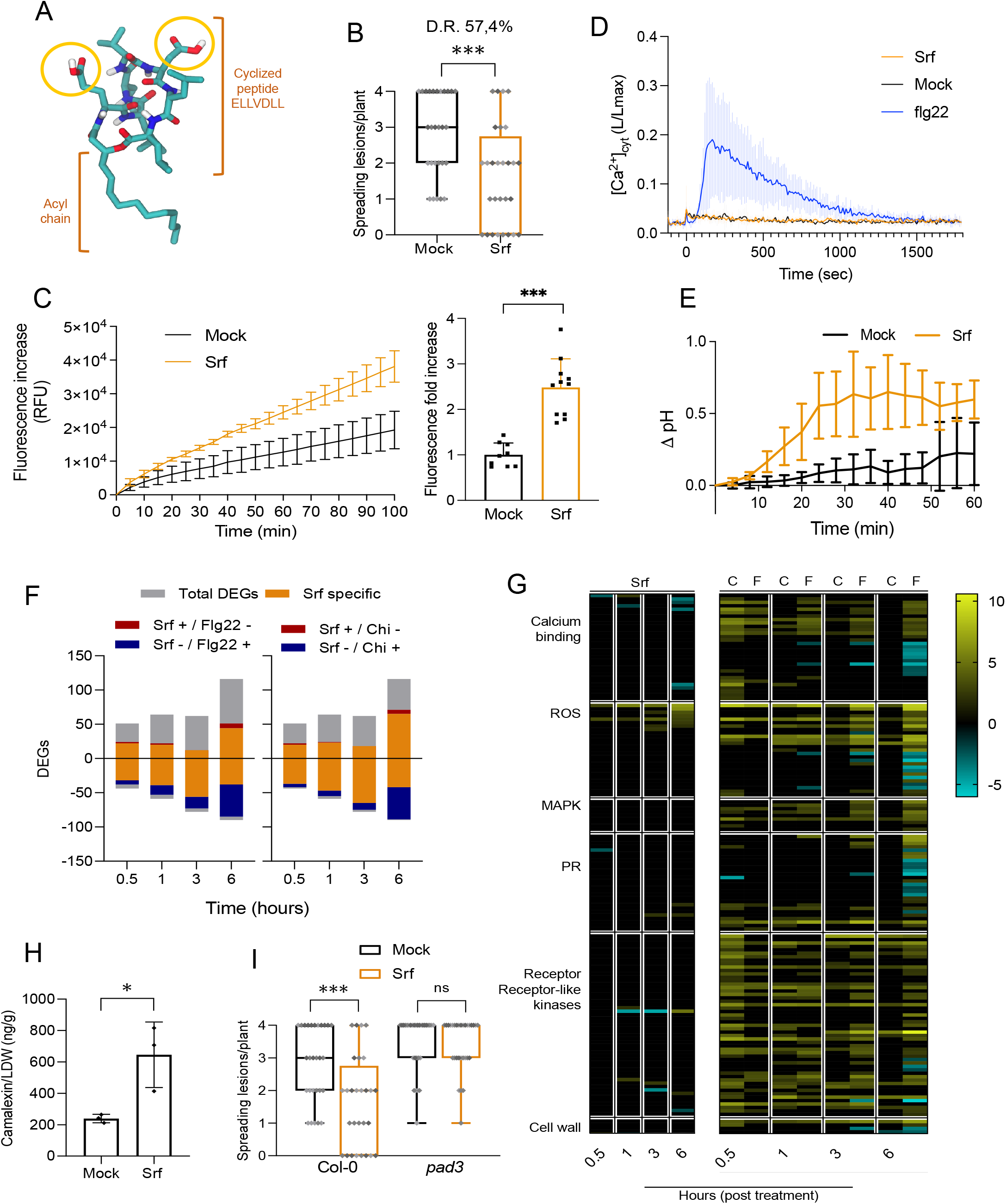
Surfactin triggers systemic resistance in *Arabidopsis* associated with atypical immune responses. **(A)** Structural model of the heptapeptide Srf (C14 acyl chain homologue) in water (Gromacs v.4.5.4). Red: oxygen, white: hydrogen, dark blue: nitrogen, light blue: carbon. The polar amino acids are circled in yellow, and other amino acids and the acyl chain constitute the non-polar part of the molecule. **(B)** Disease incidence caused by *Botrytis cinerea* in *Arabidopsis* Col-0 plants pre-treated with Srf (10 μM) or not (mock treatment, 0.1 % ethanol) (n=28 replicates from three independent experiments). The box plots encompass the 1st and 3rd quartiles, the horizontal line indicates the median, and bars extend from the lower to the higher values. Disease reduction (D.R.) is calculated from the mean values of both treatments. Significant difference \*\*\**P*<0.001, two-tailed *t*-test. **(C)** Burst in intracellular ROS species in *Arabidopsis* Col-0 roots upon treatment with Srf (10 μM). Left, time course of [ROS]_intra_ accumulation (Relative Fluorescence Units, RFU) with data at each time point representing mean ± SD, n=3 (independent root samples). Right, fold increase in fluorescence values ± SD, at 30 min after the addition of Srf compared to mock-treated roots. Data are pooled from three independent experiments (total n=9) and asterisks indicate significant difference (\*\*\**P*<0.001, two-tailed *t*-test). **(D)** [Ca^2+^]_cyt_ kinetics in Srf-treated (10 μM) or flg22-treated root tissues (1 μM) (n=6) compared to mock treatment in the *Arabidopsis* Col-0*^AEQ^* reporter line. Results are represented as luminescence counts per second relative to total luminescence counts remaining (L/Lmax; mean ± SD). Experiments were repeated three times with similar results. **(E)** pH variation in Col-0 root medium following mock treatment or addition of 10 µM Srf. Values on the graph are normalized to pH of the first time point ± SD and are from one representative experiment (n=4) out of 2 independent experiments showing similar results. **(F)** Number of DEGs (Log_2_ Fold Change > 2, P<0.05) in *Arabidopsis* root cells determined via RNAseq for each time point in response to Srf treatment (10 μM). Our data were compared with those reported for DEGs in response to flg22 (1 μM) and chitin (Chi, 1 mg/ml)^18^ and bars are subdivided by the number of genes specifically responding to Srf and by the number of genes differentially (oppositely) regulated by Srf and the two MAMPs. **(G)** Heatmap of the expression of genes putatively associated with plant immune responses (listed in Supp table 1) that were modulated upon Srf treatment (S, left) (10 μM) and compared with their expression in response to flg22 (1

We first performed quantitative and time-resolved measurements of early responses commonly associated with MAMP perception in *Arabidopsis* and other plants. [ROS]_apo_ burst is almost invariably associated with PTI^7^ but, by contrast to treatment with the MAMP flagellin-derived peptide flg22 or with chitin, we did not observe a [ROS]_apo_ burst in *Arabidopsis* root cells treated with Srf based on a horseradish peroxidase-luminol assay (**Suppl Fig 2**). Srf-mediated IR against *B. cinerea* is fully conserved in the *rbohD* mutant lacking functional plasma membrane NADPH oxidase RBOHD responsible for MAMP-induced [ROS]_apo_ burst^13^ (**Suppl Fig 3**). Hence, Srf-mediated activation of IR in the root does not require RBOHD^14^. However, Srf triggered a fast and consistent increase in intracellular ROS ([ROS]_intra_) in root loaded with the fluorescent probe DCFH-DA (**Fig 1C**). This Srf-triggered [ROS]_intra_ burst is also observed in the *rbohD* mutant (**Suppl Fig 4**), suggesting it is not caused by the uptake of apoplastic ROS via aquaporins (see **Suppl Fig 5** for response to flg22) but may originate from different organelles as reported for abiotic stresses or other small microbial compounds^7, 15–17^. Calcium influx typically associated with PTI in plants^8^ was tested upon elicitation by Srf using an aequorin-based bioluminescence assay. It did not reveal any significant Ca^2+^ increase ([Ca^2+^]_cyt_) in root of the Col-0*^AEQ^* reporter line in contrast to the increase observed upon flg22 treatment (**Fig 1D**) or in response to chitin (**Suppl Fig 6**). On the other hand, medium alkalinization occurs within minutes after Srf treatment (**Fig 1E**), which indicates H^+^/K^+^ exchange possibly leading to membrane depolarization^9^. However, no significant increase in conductivity was measured in the medium following Srf treatment (**Suppl Fig 7**) indicating that the lipopeptide does not affect plasma membrane (PM) integrity and does not cause massive electrolyte leakage. Cell viability tests confirmed that Srf is not toxic for *Arabidopsis* root cells at concentrations up to 50 µM (**Suppl Fig 8**).

Next, we explored early changes in the root transcriptome profile induced by Srf via time course RNAseq analysis (30 min, 1h, 3h and 6h post treatment) using the same setup previously reported for flg22 and the fungal MAMP chitin^18^. Data revealed a relatively low transcriptional response to Srf elicitation over all sampling times with a total of 564 differentially expressed genes (DEGs, Log_2_ Fold Change > 2, p<0.05; **Fig 1F**) compared to approximately 5000 DEGs and 2000 DEGs reported upon flg22 and chitin treatment respectively^18^. While MAMPs mainly up-regulate early responsive genes (30 min – 1 h)^18, 19^, an almost equal number of up- and down-regulated DEGs were observed upon Srf treatment at all time points (**Fig 1F**), with about half of the transcriptional changes specific to Srf elicitation (47,9% and 58% compared with flg22 and chitin respectively)^18^. Strikingly, many of the Srf down-regulated genes are upregulated by flg22 and chitin^18^ (**Fig 1F, Suppl Table 1**). Differential expression was confirmed by quantitative RT-PCR performed on some selected genes in plantlets elicited with the lipopeptide and with chitin (**Suppl Fig 9**). More specifically, the expression of genes typically associated with early immune signaling (receptor-like kinases, [ROS]_apo_ burst, calcium signaling or MAPK phosphorylation cascade^10^) or defense mechanisms (pathogenesis-related (PR) proteins, callose deposition, lignification) is not modulated or down-regulated by Srf by contrast with MAMP treatment (**Fig 1G, Suppl Table 1**).. However, *CYP71A12*, encoding a key enzyme of the camalexin biosynthesis pathway^20^, is among the late-responsive genes (6h) strongly stimulated by Srf. In accordance, we measured significantly higher amounts of this phytoalexin, which is toxic to *B. cinerea*^21, 22^, in infected leaves of Srf-treated plants compared with mock treatment (**Fig 1H**). The key role of camalexin in disease control was confirmed by the loss of Srf-triggered resistance in the *pad3* mutant^23^ unable to form camalexin^22^ (**Fig 1I**). Thus, by contrast to PTI which is associated with substantial transcriptional reprogramming^19^, immunity stimulation by Srf does not lead to major changes in the expression of genes involved in signaling and defense.

Since the molecular basis of Srf-induced immune activation is signal-specific, we hypothesized that plant cells perceive lipopeptides by a mechanism that differs from pattern sensing. Srf possesses both a peptidic moiety and a fatty acid tail, but its IR-eliciting potential is fully conserved in *Arabidopsis* mutants lacking functional PRRs that recognize either bacterial proteinaceous immunogenic patterns or acyl chain epitopes such as medium chain 3-hydroxy fatty acids and HAAs^24, 25^ (**Suppl Fig 10**). Srf elicitation is not significantly affected either in mutants lacking co-receptors required for proper functioning of a wider range of PRRs detecting immunogenic peptides such as Pep1^26^, nlp20^27^ and IF1^28^ nor in the *bik1 pbl1* double mutant lacking RLCKs that act downstream of the PRR-co-receptor complexes (**Suppl Fig 10**). Although we only tested a small subset of the multitude of PRRs potentially expressed in *Arabidopsis*^6^ and although early cellular signaling may be BIK1/PBL1-independent^29^, our data strongly suggest that *Arabidopsis* does not sense Srf via PRR-type cell surface sentinels. This is in accordance with previous data from tobacco, which showed that Srf is still active on protease-treated cells and that there is no refractory state upon repeated Srf treatment unlike typically observed for PTI^30^.

Due to their amphipathicity, CLPs readily interact with biological membranes, causing pore formation and membrane disruption responsible for their antimicrobial activities^31^. Such an adverse effect is not expected on plant membranes, but we hypothesized that Srf perception by root cells might primarily rely on its interaction with the lipid phase of the PM. Complex sphingolipids glucosylceramides (GluCer) and glycosyl inositol phosphorylceramides (GIPC) constitute more than 30% of *Arabidopsis* PM lipids and are key components required for membrane integrity and functionality, notably by forming ordered nano-domains with sterols^32–34^. *In silico* docking simulation first revealed a more favorable interaction of Srf with GluCer or GIPCs than with the other typical plant PM lipids PLPC (1-Palmitoyl-2-linoleoyl-sn-glycero-3-phosphocholine as phospholipid) and ß-sitosterol (as main sterol) (**Fig 2A**). To test this experimentally, we generated biomimetic liposomes using commercially available GluCer, PLPC and ß-sitosterol. Isothermal titration calorimetry performed on liposomes with increasing composition complexity in such lipids showed the highest binding affinity of Srf to model membranes containing GluCer (**Fig 2B**). In support of a preferential interaction with sphingolipids, molecular dynamic (MD) simulation on the same ternary lipid system showed the specific insertion of Srf in the vicinity of GluCer molecules or in GluCer-enriched areas in the membrane (**Fig 2C**). In light of these results, we tested Srf elicitor activity on the *Arabidopsis* ceramide synthase mutant *loh1* (LONGEVITY ASSURANCE 1 HOMOLOG1) which is depleted in these complex sphingolipids^35, 36^. We observed strongly reduced [ROS]_intra_ responses (**Fig 2D**) as well as loss of IR to *B. cinerea* infection in *loh1* compared to wild-type plants (**Fig 2E**). Such lipid-dependent [ROS]_intra_ elicitation was also observed for other IR-eliciting CLPs such as orfamide and WLIP^2^ isolated from beneficial pseudomonads that resemble Srf in size and amphiphilic character (**Suppl Fig 11**). The CLP immunogenic activity thus relies on an intricate interaction with PM sphingolipids as reported for other microbial compounds^36–38^.

**Fig. 2.**
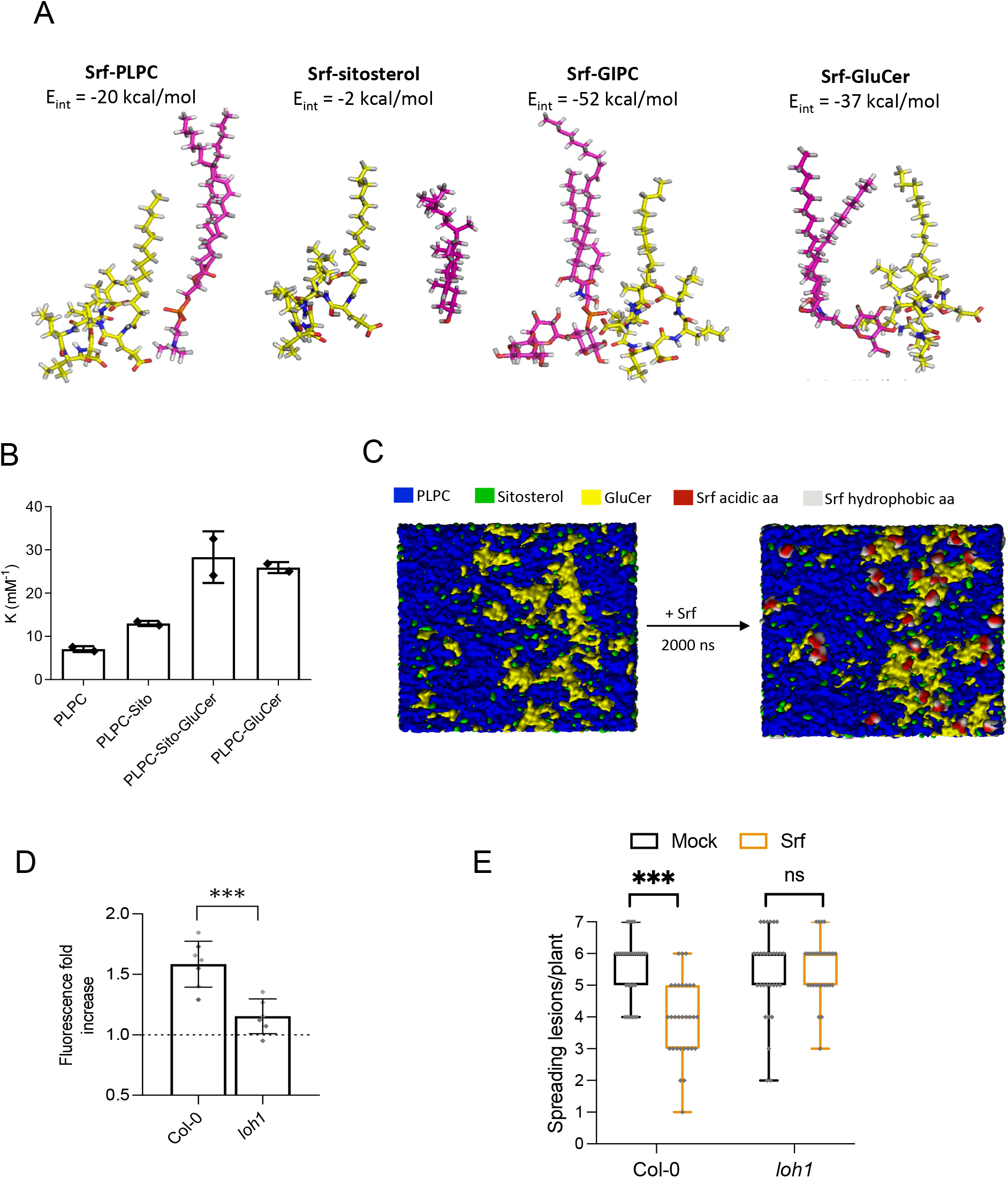
Affinity for sphingolipids determines CLP-triggered immunity. **(A)** *In silico* docking simulation of the interaction between Srf and plant PM lipids with their associated energy of interaction (E_int_). A lower E_int_ value indicates a more favorable interaction. Hydrogen, oxygen and phosphate atoms are respectively represented in grey, red and blue. Carbon atoms of Srf are in yellow and carbon atoms of GluCer, Sito and PLPC are in pink. **(B)** Binding coefficient (K) of Srf to liposomes with different lipid compositions. Graph presents values from two independent experiments, mean ± SD. **(C)** Molecular dynamics simulation of Srf insertion in GluCer-enriched domains of a PLPC-Sito-GluCer bilayer. Left: Top views of bilayers before and after Srf insertion (right). **(D)** [ROS]_intra_ accumulation in roots of Col-0 and *loh1* mutant. Data represents fold increase in fluorescence values ± SD (n=6 from two independent experiments) at 30 min after Srf addition (10 μM) or not. Significant difference \*\*\**P*<0.001, two-tailed *t*-test. **(E)** Disease incidence of *B. cinerea* in *Arabidopsis* Col-0 and *loh1* mutant plants, pre-treated with Srf (10 μM) compared with mock treatment (n=30 from two independent experiments). Data are represented as in fig 1B. ns = not significant, \*\*\**P*<0.001, two-way ANOVA and Sidak’s multiple-comparison post-test.

By inserting into lipid bilayers, Srf may transiently affect the local structure of membranes. Indeed, neutron reflectivity (NR) experiments (see **Suppl Fig 12** for deuterated Srf synthesis and characterization) demonstrate that Srf exclusively inserts into the outer leaflet of PLPC-ß-sitosterol-GluCer model membranes (**Fig 3A**). This is supported by MD simulation showing that the Srf peptide backbone preferentially positions at the level of the polar lipid heads of the membrane (**Suppl Fig 13**). Srf insertion does not affect the lipid chain-chain interaction as shown by WAXS and FTIR (**Suppl Fig 13**). In addition, NR data indicated that Srf insertion results in a decrease in membrane thickness (from 40 to 36Å), which is more pronounced in ternary membranes than in membranes lacking GluCer (from 43 to 41Å) (**Suppl Table 4**). Analysis of the nanoscale morphology of supported PLPC-ß-sitosterol-GluCer bilayers by Atomic Force Microscopy confirmed this membrane thinning caused by Srf insertion (**Suppl Fig 14**). An additional impact of the lipopeptide on PM physical properties was derived from coarse-grained MD simulation which revealed a strong curvature-inducing effect mediated by Srf docking on ternary membranes (**Fig 3B**).

**Fig. 3.**
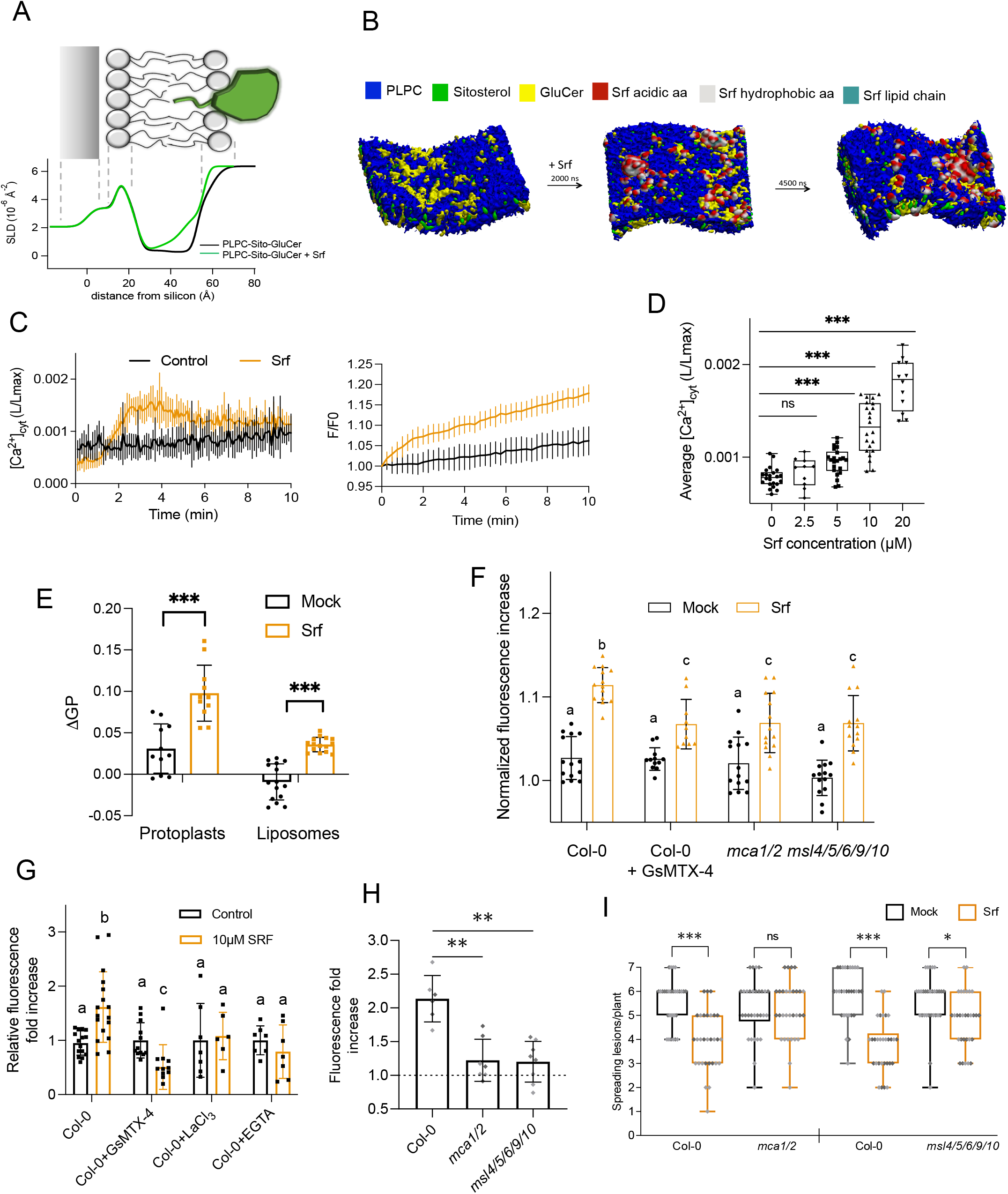
Srf causes membrane deformation and activates mechanosensitive channel-dependent immune responses. **(A)** Membrane thickness determined via neutron scattering length density (SLD) profiles of supported PLPC-Sito-GluCer membrane before (black) and after (green) Srf addition to the final 95:5 membrane: Srf molar proportion (0.24 μM) (below). Illustration (above) presents the correspondence between regions in the SLD profile and specific zones in the membrane. **(B)** Molecular dynamics simulation of Srf-induced membrane curvature. **(C)** Left, [Ca^2+^]_cyt_ kinetics in Srf-treated (10 μM) root cell protoplasts (n=6) compared to mock treatment in the *Arabidopsis* Col-0*^AEQ^* reporter line. Results are represented as luminescence counts per second relative to total luminescence counts remaining (L/Lmax; mean ± SD). Experiments were repeated three times with similar results (see Suppl Fig 16 for additional experiments). Right, increase in [Ca^2+^]_cyt_ detected upon loading root protoplasts of Col-0 with Fluo-4 in mock- or Srf-treated (10 μM). Experiments were repeated three times with similar results. **(D)** Dose-dependent [Ca^2+^]_cyt_ increase induced by Srf in root protoplasts of *Arabidopsis* Col-0^AEQ^. Values are the average of L/Lmax values from 1.5 to 4 min after treatment corresponding to the top of the peak. Mean ± SD of at least 10 technical replicates from at least five independent experiments. Asterisks indicate statistically significant differences to the mock treatment (ns= no significant difference; \**P*<0.05; \*\*\**P* < 0.001; (a) two-tailed *t*-test; (b) Welch and Brown-Forsythe ANOVA). **(E)** Change of laurdan generalized polarization (ΔGP) in Srf-treated (10 μM) Col-0 root protoplasts and in liposomes reflecting a change of membrane rigidity. ΔGP is defined as the subtraction of GP measured at 10 min following treatment and GP measured before treatment. Mean ± SD of 12 (for protoplasts) and 15 (for liposomes) replicates from 8 (for protoplasts) and 5 (for liposomes) independent experiments. \*\*\**P*<0.001, two-way ANOVA and Sidak’s multiple comparison test. **(F)** [Ca^2+^]_cyt_ response measured with Fluo-4 upon Srf elicitation (10 µM) in *Arabidopsis* Col-0 root protoplasts with and without pre-treatment with the mechanosensitive channel blocker GsMTX-4 (10 min incubation, 7.5 µM)(n=10) and in root protoplasts of the *mca1/2*, and *msl4/5/6/9/10* mutants (n=14). Mean ± SD from four independent experiments. Letters represent statistically different groups at α = 0.05 (two-way ANOVA and Tukey’s multiple-comparison post-test). **(G)** [ROS]_intra_ accumulation upon addition of 10 μM Srf to *Arabidopsis* Col-0 roots upon pre-treatment or not (Col-0, n=16) with the mechanosensitive channel blocker GsMTX-4 (10 min incubation, 7.5 µM) (n=12), with the non-selective Ca^2+^ channel blocker LaCl_3_ (10 mM) (n=7) and the Ca^2+^ chelator EGTA (1 mM) (n=7). Data represent fold increase in fluorescence values 30 min after Srf addition compared to mock-treated roots. Mean ± SD calculated from data from two independent experiments. Letters represent statistically different groups at α = 0.05 (two-way ANOVA and Tukey’s multiple-comparison post-test). **(H)** [ROS]_intra_ accumulation in *Arabidopsis* Col-0 (n=6), *mca1/2* (n=7), and *msl4/5/6/9/10* (n=8) roots following Srf treatment (10 μM). Data represent fold increase in fluorescence values 30 min after Srf addition compared to mock-treated roots. Mean ± SD from two independent experiments. \*\**P*<0.01, two-tailed *t*-test. **(I)** Disease incidence of *B. cinerea* in *Arabidopsis* Col-0, *mca1/2*, and *msl4/5/6/9/10* mutant plants, mock- or Srf pre-treated (10 μM)(each n=30 from two independent experiments represented as differently shaded grey values). Data are represented as in fig 1B.

In light of these biophysical data, a clear impact of Srf on PM structure can be predicted but in integral root cells, the PM is physically connected to the thick and mechanically strong cell wall polymer matrix, which provides structural support and might stabilize the membrane into a flat conformation under low tension^39^. We thus next tested early Srf-induced immune responses in cell wall-free protoplasts. Use of PPs renders the PM more susceptible to deformation, which was used to study responses to cell swelling or shrinkage/expansion during osmotic stresses^40, 41^. As in root cells, Srf triggered a consistent [ROS]_intra_ burst in freshly isolated protoplasts (**Suppl Fig 15**). However, in contrast to roots, a significant calcium influx, was observed in Srf-treated protoplasts by using the Col-0*^AEQ^* reporter line and also by loading Col-0 with the Fluo4-AM probe (**Fig3C and Suppl Fig 16**). This Srf-induced Ca^2+^ influx is comparable in amplitude to the one induced by MAMPs (**Suppl Fig 17**). It involves some PM channels since it is abolished in protoplasts pre-treated with the general channel blocker LaCl_3_ (**Suppl Fig 18**) and considering that the lipopeptide does not cause any detrimental effect on protoplast viability at the concentration used (**Suppl Fig 19**). Additional assays on protoplasts revealed that activation of early responses by Srf requires threshold concentrations of 5-10 µM both for calcium influx (**Fig 3D**) and [ROS]_intra_ burst (**Suppl Fig 20**), which is much higher than MAMPs detected at nanomolar concentrations. This further indicates that Srf perception is not mediated by a high-affinity receptor-based detection system and is in accordance with the mechanism predicted from biophysics in which threshold amounts of Srf molecules must dock into sphingolipid domains in order to modulate PM structure. We tested the impact of the lipopeptide on protoplast membrane fluidity via measurements of laurdan generalized polarization (laurdan GP) related to the lipid bilayer order. Our results show that Srf treatment led to a significant increase in ΔGP values indicating a clear membrane rigidification effect as also observed upon interaction of the lipopeptide with PM mimicking liposomes (**Fig 3E**).

Altogether, these data obtained with protoplasts support the relevance of PM deformation in the response to Srf. We therefore hypothesized that insertion of the CLP could induce physical constrains resulting in increased lateral tension sufficient for activating mechano-sensitive (MS) ion channels, in a process similar to the one observed for some anionic amphipathic chemicals^42, 43^. This was supported by the reduced calcium influx observed upon pre-treatment of Col-0*^AEQ^* protoplasts with the specific MS channel blocker GsMTX-4 (**Suppl Fig 21**). Among stretch sensitive mechanosensors identified so far in plant cells, MSL9, MSL10 and MCA1/2 localize in the PM ^44–46^ but do not require RLCK-mediated phosphorylation of the cytoplasmic domains for gating unlike other MS ion channels such as OSCA1.3 which needs BIK1 phosphorylation to be activated ^47^. Using Fluo-4, we thus tested protoplasts prepared from the quintuple *msl4/5/6/9/10*^48^ and the double *mca1/2*^49^ mutants for their response to Srf and observed a significantly decreased calcium influx, to the same extent as chemical inactivation with GsMTX-4 in Col-0 (**Fig 3F**).

We next evaluated the effect of inactivation or knock-out of *msl* and *mca* channels on intracellular ROS burst as early response of root tissues elicited by Srf. Pre-treatment with GsMTX-4, LaCl_3_ or with the Ca^2+^ chelator EGTA eliminated the ROS burst triggered by Srf in root tissues (**Fig 3G**), supporting the importance of MS channels in the response and indicating that ion fluxes acts upstream of or are interdependent of [ROS]_intra_^8^. An almost complete loss of [ROS]_intra_ burst was also observed upon Srf treatment in the *msl4/5/6/9/10* and *mca1/2* mutants as compared to Col-0 (**Fig 3H**). In addition, *mca1/2* and *msl4/5/6/9/10* plants were strongly impaired in mounting systemic resistance against *B. cinerea* upon Srf treatment (**Fig 3I**), further indicating that functional MS channels are necessary for full response of *Arabidopsis* to Srf elicitation on roots. Data on protoplasts show that Srf may trigger some calcium transients as early immune-related event but a detectable Ca^2+^ influx is not required for defense activation in plantlets, which correlates with the fact that no downstream components of calcium signaling are up-regulated upon perception of the lipopeptide.

Collectively, although contributions of other channels cannot be ruled out^47^, our data provides evidence for a key role of PM-located mechanosensors in lipopeptide-induced plant defenses. The relative contribution of each channel remains to be determined as they display specific properties in terms of sensitivity to membrane tension and ion selectivity. MCA1/2 are described as genuine transporters of Ca^2+^ ^46, 50^ while MSL10 is regarded as a non-selective ion transporter that is indirectly involved in calcium signaling upon wounding^51^ and response to hypo-osmotic shock in cell swelling^41^. Both channels may thus act in a coordinated fashion to tailor ion fluxes leading to cellular responses and PM depolarization. As previously reported for other plant species^2^, treatment with Srf prepares *Arabidopsis* to mount defense responses culminating in the systemically expressed IR phenotype. We provide new insights into the molecular basis of the well-known long-standing process of CLP-triggered plant immunity activation by unveiling a new lipid-mediated mechanism for the detection of these molecules at the cell surface. We infer from our data that CLP insertion into sphingolipid-enriched PM domains causes deformation and increases lateral tension in the membrane leading to rearrangement of the MS protein complexes and gating of the channels. This allows ion influx and initiates chemical signaling that can be integrated by root cells to activate early immune responses in a process that remains to be deciphered. Such a lipid-dependent perception at the cell surface may apply also to other bacterial amphiphilic IR elicitors such as acyl-homoserine lactones and rhamnolipids which also readily interact with membrane lipids and may thus be perceived via similar mechanisms^2, 52–55^. The nature of PM lipids widely varies across plant species^34^. This could explain, at least in the case of Srf, why this molecule triggers immunity in dicots but is not very active on monocots^2^. We assume that the effect of a CLP on a particular target membrane is also fine-tuned by precise structural traits in the molecule. It may explain why some CLPs produced by *Pseudomonas* leaf pathogens act as virulence factors in a wide range of plants by causing necrosis via pore formation in cellular membranes^56^. However, further investigation is required to capture the physico-chemical rules governing lipid selectivity and CLP insertion dynamics.

As the two components of the plant immune system, PTI works in concert with effector-triggered immunity (ETI) for mounting robust defense responses to biotrophic invaders but ETI is not efficient against necrotrophic pathogens^5, 57^. Here, we describe a novel molecular mechanism of defense activation in plants, which provides resistance to the necrotroph *B. cinerea* via a unique process not related to the receptor-based surveillance system involved in the recognition of MAMPs by plant cells or in the perception of the Pam_3_CSK_4_ analog of triacylated lipopeptides produced by *Staphylococcus aureus* and acting as agonists of Toll Like-type PRRs in metazoans^58^. It therefore provides new insights in plant-microbe interactions mediated by small chemicals from beneficial bacteria. Collectively, our data show that Srf perception leads to specific immune activation signature regarding the type, timing, and amplitude of early defense-related events and the weak transcriptional reprogramming as compared to PTI. This may explain why elicitation by Srf is cost-effective for the host plant as it does not result in growth-defense trade-off^59, 60^ nor does it cause a strong response associated with the alertness state or a hypersensitive reaction leading to cell death. Using CLPs as elicitors would enable bacteria to bypass a strong immune response and avoid their rejection as undesirable associate. Further investigations are needed for a comprehensive understanding of the whole process from perception to systemic signaling but the mechanistic basis of CLP-induced plant resistance reported here should contribute to rationally implement the use of these compounds or their producers as bio-sourced alternatives to chemicals in sustainable agriculture. 1μM) and chitin (1 mg/ml) (F and C respectively, right) based on published data^18^. Colour scale represents Log_2_ FC (> 2, P<0.05). **(H)** Camalexin response associated with IR triggered by Srf. Camalexin accumulation 96 hours post *B. cinerea* inoculation (hpi) in *Arabidopsis* Col-0 leaves of mock- or 10 µM Srf-treated plants at the root level. Graph shows values obtained in one experiment with each value representing a sample of five plants pooled together. Asterisks indicate significant difference with ns, not significant; \**P*<0.05; \*\*\**P*<0.001; two-tailed *t*-test. **(I)** Disease incidence of *B. cinerea* in *pad3* mutant pretreated with 10 µM Srf or mock-treated at the root level (n=30, values obtained from three independent experiments, presented as differently shaded grey values). Data are represented as in fig 1B. Asterisks indicate significant difference with ns, not significant; \**P*<0.05; \*\*\**P*<0.001; two-way ANOVA and Sidak’s multiple-comparison post-test.

## Acknowledgments

We are grateful to Prof. Ivo Feussner at University of Goettingen for providing us with *Arabidopsis* lipid mutants. We thank Aurélien Legras, Sébastien Steels, Catherine Helmus, and Adiilah Mamode-Cassim for their excellent technical support.

## Author contributions

J.P., G.G., M.D. and M.O. conceived and designed experiments; J.P., G.G., H.I., A.A., V.R., W.P.L-L, E.D., M.N.N., M.M-G., S.E., A.K., P.B. and Y.F.D. performed experiments and analyzed data; J-M.C., M.G. and L.L. performed modeling; S.D., B.D.C, S.R., T.N., S.R., M.H. and C.Z. substantially revised the manuscript and were involved in the discussion of the work; M.D. and M.O. supervised the study and provided funding.

