## Supplementary data for "Mechanosensing and Sphingolipid-Docking Mediate Lipopeptide-Induced Immunity in *Arabidopsis*"

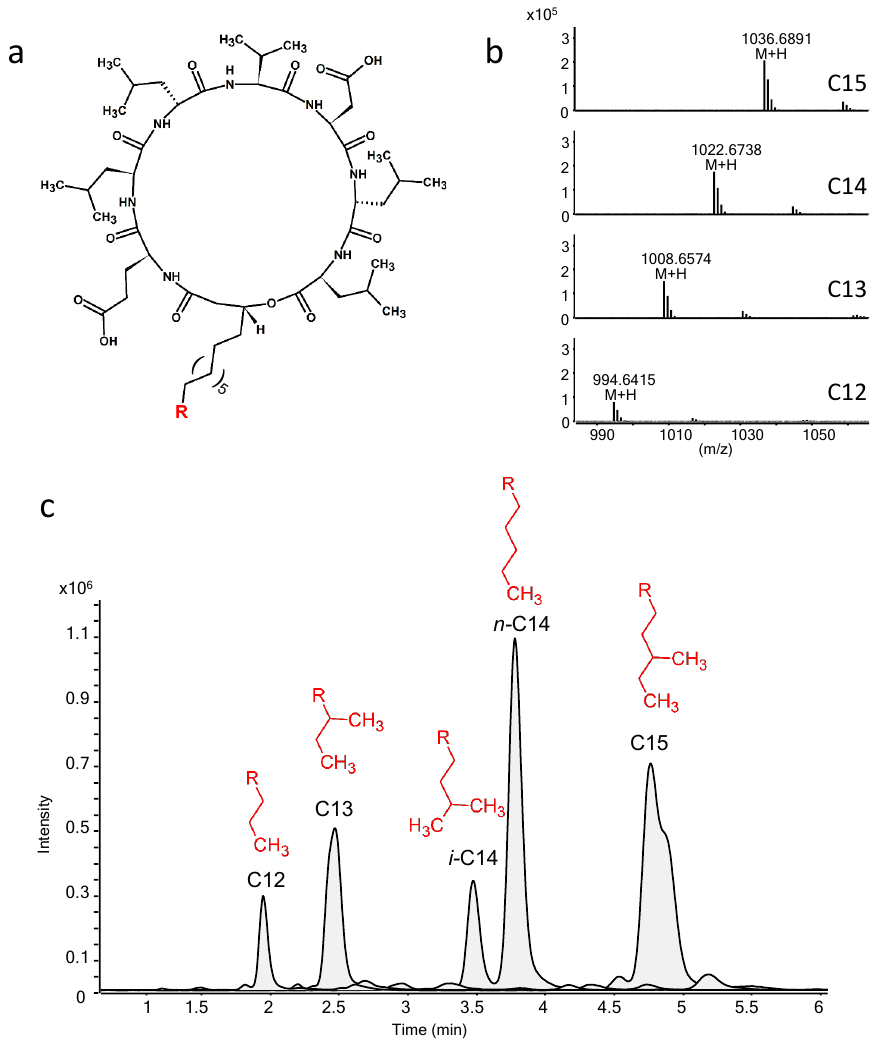

Suppl Fig 1. *Bacillus velezensis* produces surfactin as a mixture of structural variants differing in length and branching type of the fatty acid chain (R in the structure presented in (a)) that are identified on the basis of the exact mass of their molecular ion (b) upon UPLC-MS profiling (c). This diversity is due to the low selectivity of the first C-domain of the multi-modular enzymatic machinery responsible for the synthesis of the compound. It allows using diverse fatty acids from the intracellular pool for binding to the first amino acid of the nascent peptide.

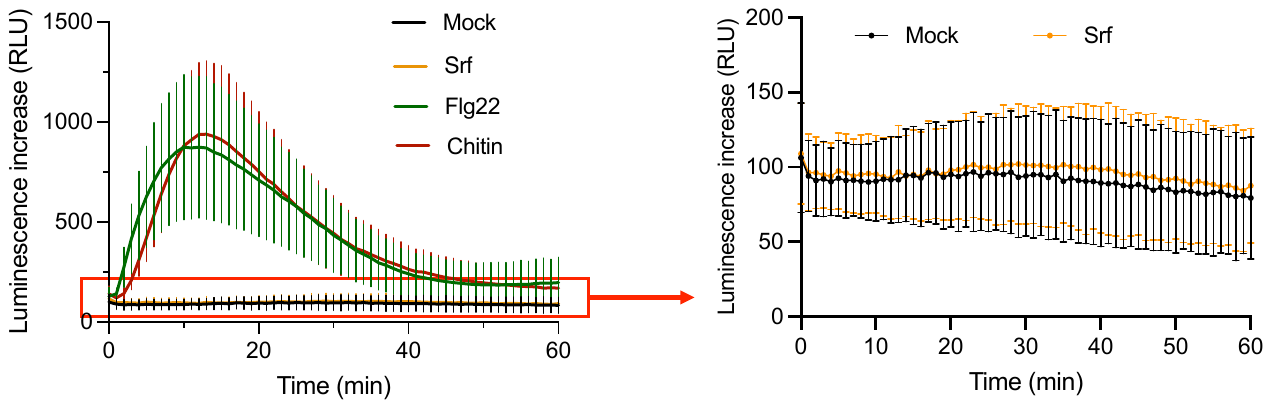

**Suppl Fig 2.** Kinetics of [ROS]_apo_ burst measured in relative luminescence units (RLUs) in *Arabidopsis* roots upon perception of 1µM flg22, 100 µg/mL chitin and 10 µM Srf. Luminescence increase in plotted by dividing the measurement at each time point by the measurement at time 0 for each treatment. Means and SD were calculated from data obtained in two independent experiments each n=8.

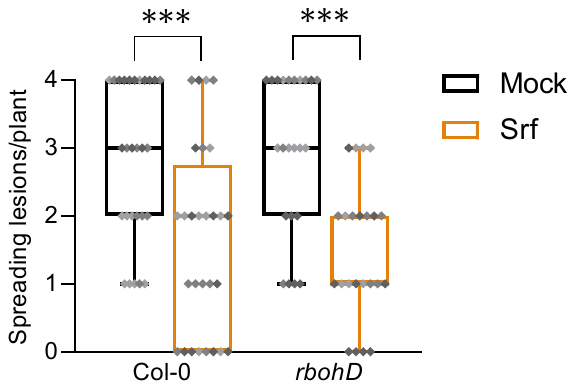

**Suppl Fig 3.** *Botrytis cinerea* disease incidence in *Arabidopsis* Col-0 and *rbohD* mutant plants, mock or Srf pre-treated (10 μM) (n=28 for Col-0 and n=23 for *rbohD*, both from three independent experiments). The box plots encompass the 1st and 3rd quartile, the whiskers extend to the minimum and maximum points, and the midline indicates the median. Asterisks indicate significant difference (****P*<0.001, two-way ANOVA and Sidak’s multiple-comparison post-hoc test).

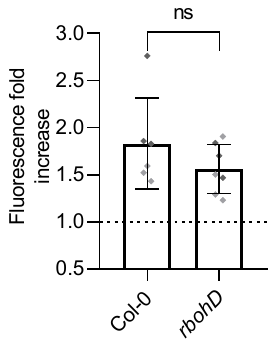

**Suppl Fig 4.** [ROS]_intra_ accumulation in Col-0 and *rbohD* roots following Srf (10 μM) treatment. Values represents fold increase in fluorescence values ± SD, 30 mins after the addition of Srf compared to mock-treated roots. Data are from two independent experiments (each n=3 or n=4) with differently shaded grey values of the symbols. ns indicates that there is no significant difference (two-tailed *t*-test).

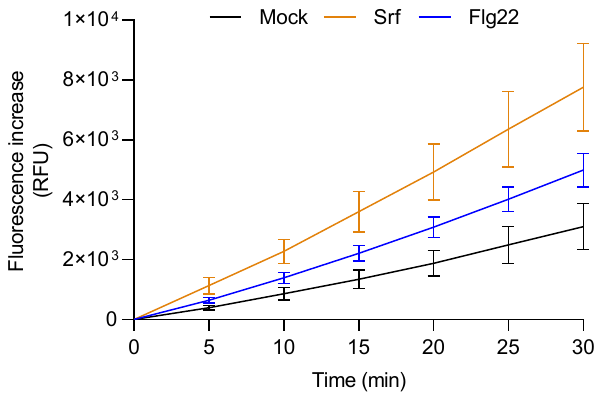

**Suppl Fig 5.** [ROS]_intra_ accumulation in *Arabidopsis* Col-0 roots following treatment with 10 µM Srf, 1 µM flg22 or mock-treated measured with the fluorescent probe DCFH-DA. Data represents one (n=4) out of 2 independent experiments showing similar results.

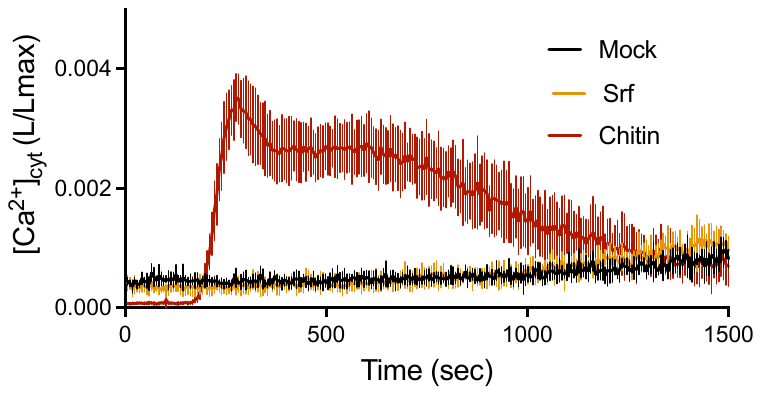

**Suppl Fig 6.** Kinetics of [Ca^2+^]_cyt_ production in *Arabidopsis* Col-0^AEQ^ roots treated with 10 µM Srf compared to 100 µg/mL chitin. Results are presented as luminescence counts per second relative to total luminescence counts remaining (L/Lmax) ± SD, n=8 from two independent experiments.

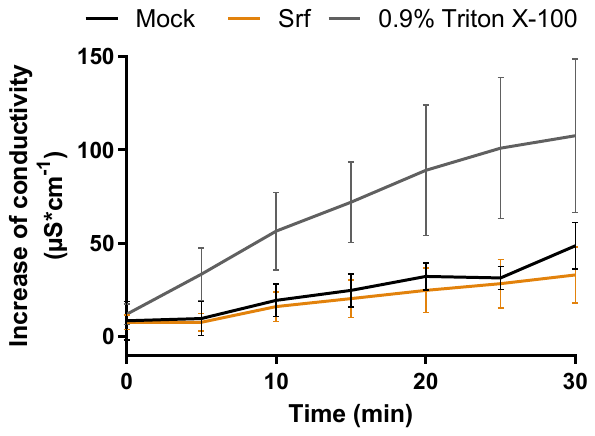

**Suppl Fig 7.** Conductivity variation in Col-0 root medium following Srf (10 µM), mock treatment or upon addition of Triton X-100 as positive control with leakage. Means and SD are from 2 independent experiments with in total n=8 for control and n=9 for Srf and Triton X-100.

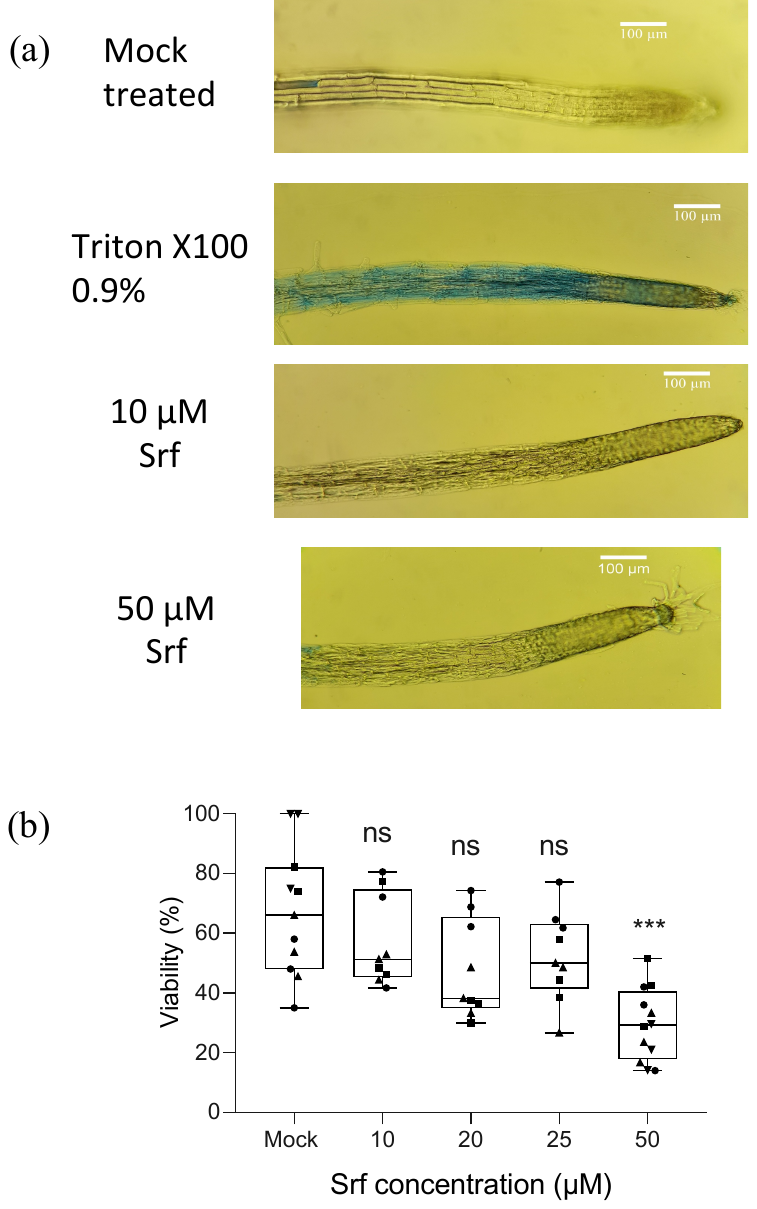

**Suppl Fig 8.** Root cell viability visualized with Evan’s blue staining following treatment with different Srf concentrations, 0.5% ethanol (negative control) or 0.9% Triton X-100 (positive control). Experiments were performed on three different individuals for each treatment with similar results.

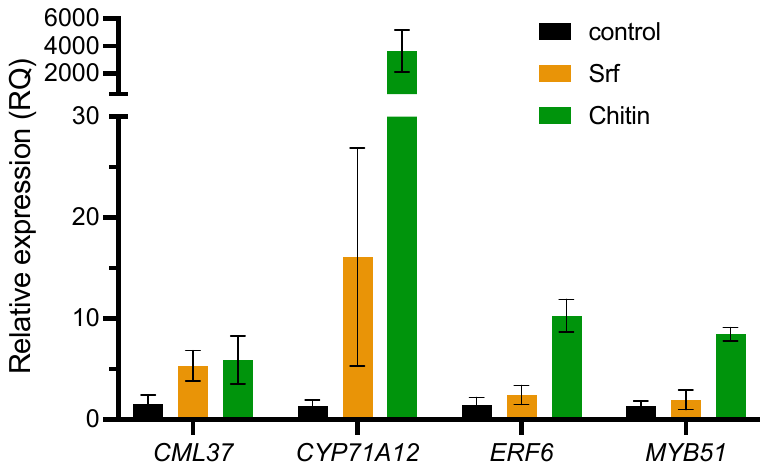

**Suppl Fig 9.** Relative expression levels of genes in roots of Col-0 plants 6 hours after treatment with surfactin (10 µM) or chitin (100 µg/mL). Expression levels are normalized to the housekeeping gene *UBQ5*. Data from one representative experiment represent mean values ± SD calculated from three biological replicates, each containing eight roots. The experiment was repeated twice with similar results.

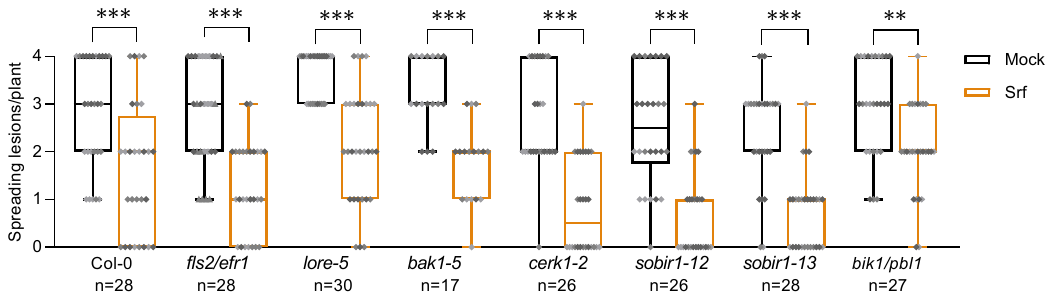

**Suppl Fig 10.** Disease incidence of *B. cinerea* in infected plants pretreated with 10 µM Srf or mock-treated at the root level in *Arabidopsis* wild-type plants (Col-0) or mutants lacking functional receptors required for the detection of either bacterial proteinaceous immunogenic patterns (*fls2/efr*^3^) or acyl chain epitopes (*lore*-5^13^), co-receptors (*bak1*-5^5^, *cerk1*-2^6^, *sobir1*-12 and *sobir1*-13^7^), or receptor-like cytoplasmic kinase (*bik1/pbl1*^4^). The box plots encompass the 1st and 3rd quartiles, the horizontal line indicates the median, and bars extend from the lower to the higher values. Asterisks indicate statistically significant differences to the mock treatment (***P*<0.01, ****P*< 0.001, two-way ANOVA and Sidak’s multiple comparison test). Data presented are from three independent experiments (n=x) presented as differently shaded grey values.

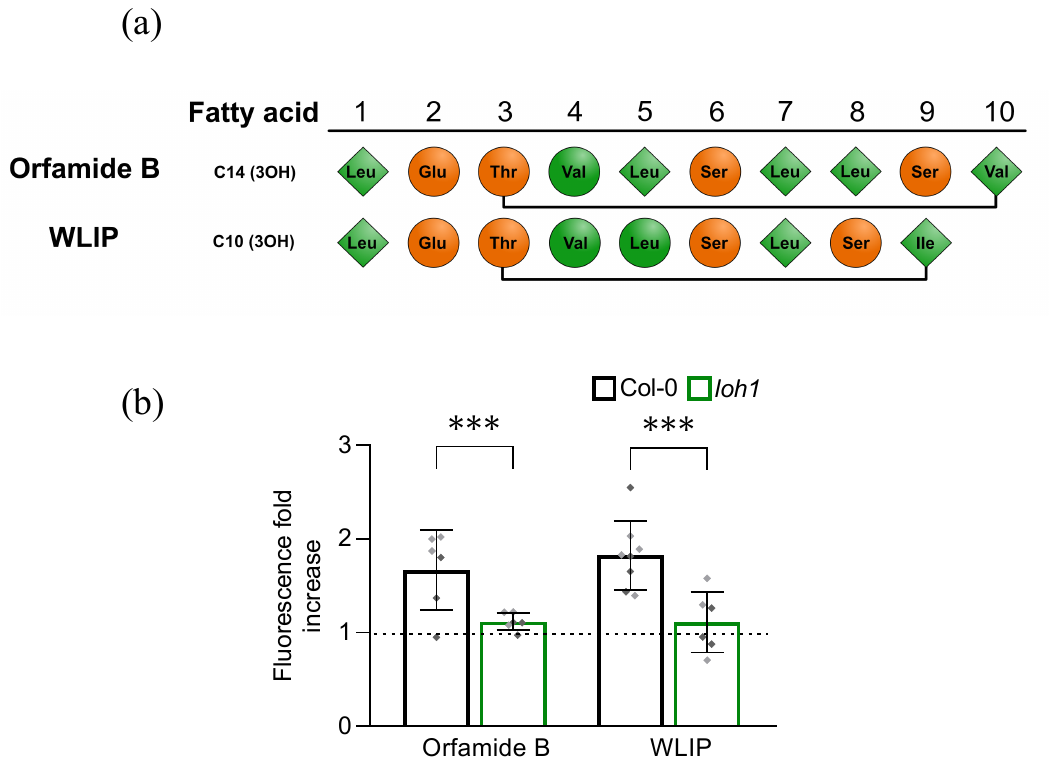

**Suppl Fig 11.** **(a)** Structures of the CLPs produced by *Pseudomonas* sp*.* used in this study. Rhombus indicates L- and circle D-amino acid; green, nonpolar; orange, polar. **(b)** [ROS]_intra_ production in roots of Col-0 (n=6) and *loh1* mutant (n=6) measured with DFCH-DA following treatment with 10 µM orfamide B or WLIP. Data represent fold increase in fluorescence values ± SD recorded at 30 min after the addition of these molecules compared to values obtained for mock-treated roots. Asterisks indicate significant difference (****P*<0.001, two-tailed t-test).

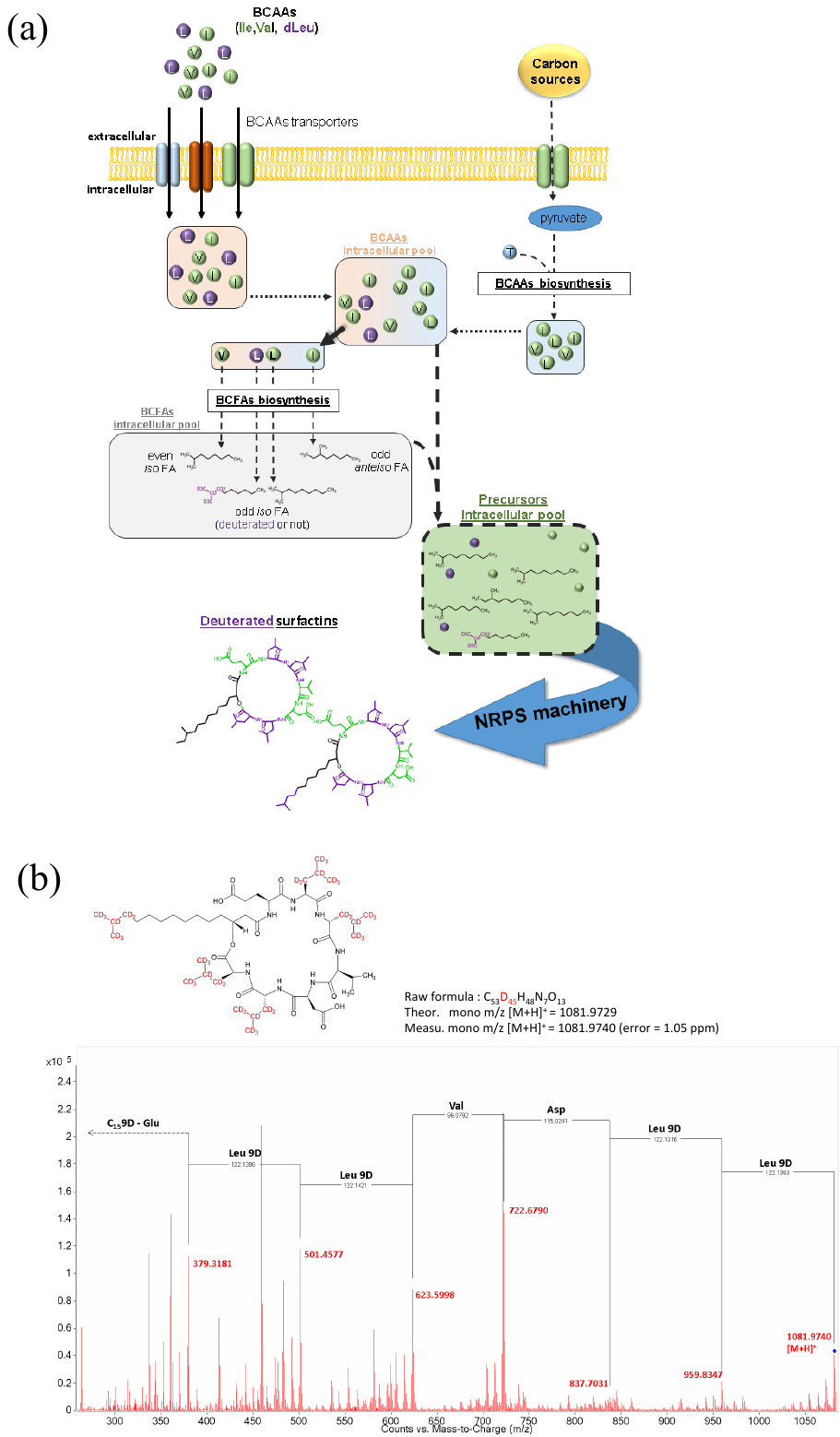

**Suppl Fig 12.** Production of deuterated Srf by exploiting the non-ribosomal enzymatic biosynthesis. **(a)** Besides the pathway using pyruvate and threonine, supplementation of the culture medium with Leu-d9 leads to enrichment of the intracellular pool in this deuterated residue via direct import involving transporters for branched-chain amino acids (BCAAs). These BCAAs are also substrates for the synthesis of branched fatty acids (BCFAs) with leucine acting as a precursor for the production of odd *iso* FAs [3] and therefore indirectly allowing incorporation of deuterium into the lipidic chain of the lipopeptide. These FAs and AAs pools, composed of a mixture of non-deuterated and deuterated residues, constitute the intracellular pool of CLP building blocks that will be activated by the multi-modular enzyme machinery in order to produce deuterated Srf variants. These variants were structurally characterized via UPLC-MS/MS based on exact mass and fragmentation as illustrated in **(b)**.

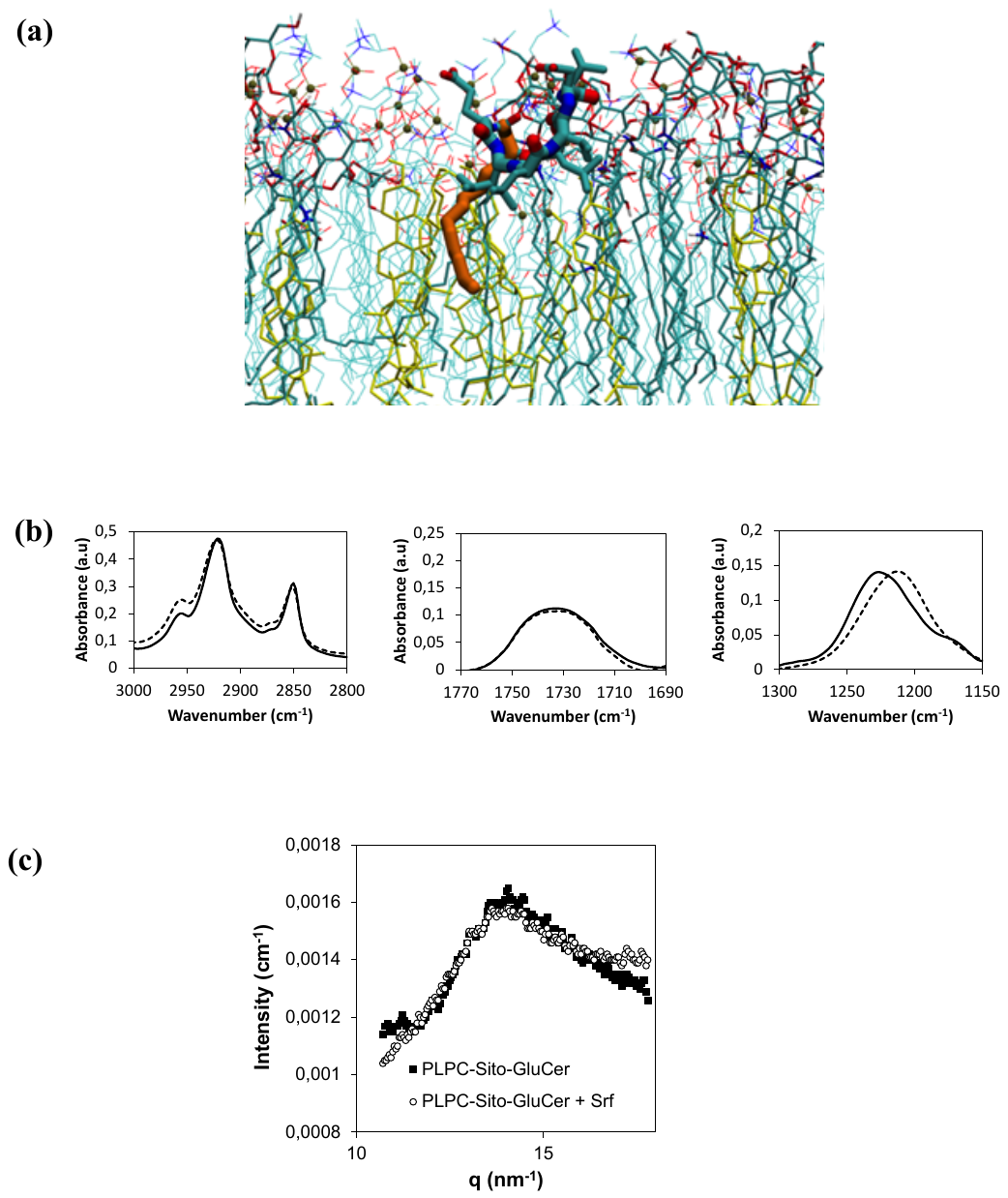

**Suppl Fig 13.** Insertion of Srf in the polar head region of the PLPC-Sito-GluCer model membrane. **(a)** Insertion of Srf (C14 acyl chain homologue) in the polar head region of the PLPC-Sito-GluCer bilayer simulated by molecular dynamics. Sito molecules are shown in yellow. PLPC and GluCer are drawn as thin lines except for the phosphorus atoms, which are represented by brown spheres. Srf is depicted as thick lines and its acyl chain is represented in orange. Light blue: carbon atoms, red: oxygen atoms, dark blue: nitrogen atoms. **(b)** Fourier-transform infrared (FTIR) spectra of multilamellar liposomes composed of ternary lipid mixture (PLPC-Sito-GluCer; full line) or of liposomes in the presence of Srf (dashed line): (left) region of alkyl chains, (middle) region of C=O ester, (right) region of P=O groups. Srf leads to the hydration of phospholipid PO^2-^ groups visualized by a shift of the band to lower wavenumbers while it has no effect on the alkyl chains and C=O ester bonds. **(c)** Wide angle X-ray scattering (WAXS) spectra. The conserved position of the typical peak obtained by WAXS for lipid bilayer, generally interpreted as a measure of the average chain-to-chain distance, demonstrates that lipid chain-chain interaction is not affected by the Srf insertion.

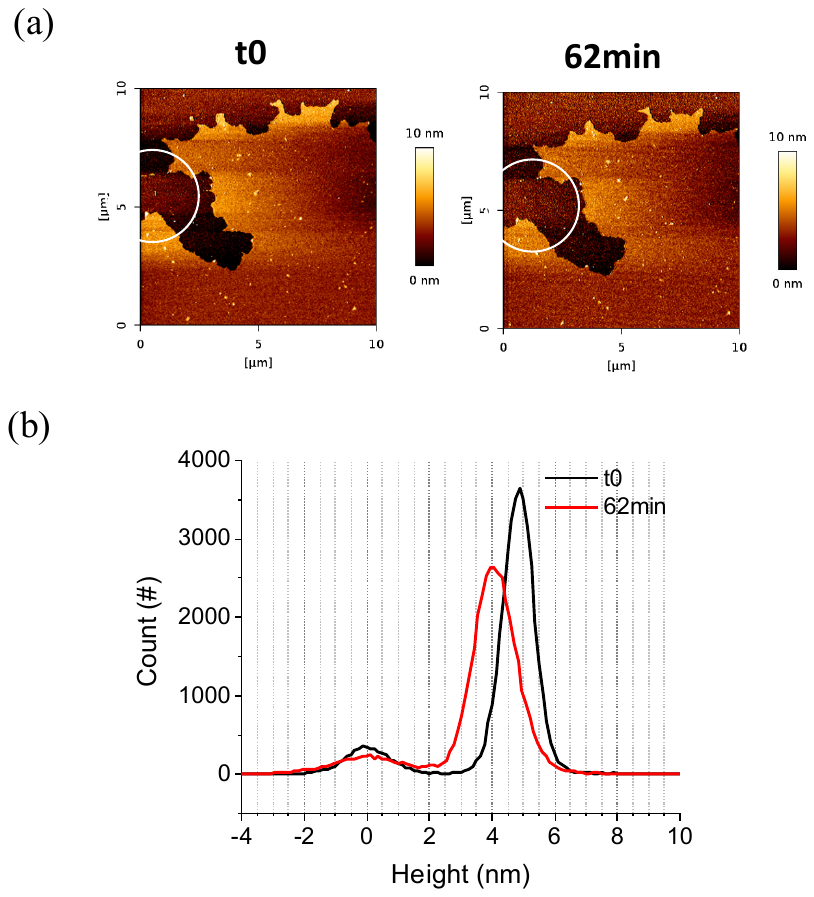

**Suppl Fig 14.** Atomic Force Microscopy confirms thickness reduction of the PLPC-Sito-GluCer bilayer following Srf insertion. **(a)** AFM topographic images before (t=0 min) and after injection of Srf (final concentration of 3 µM) (t=62 min) on a PLPC-sito-GluCer mica-supported lipid bilayer (SLB). **(b)** AFM height density profiles recorded on small areas (circled in white in (a)) of the PLPC-Sito-GluCer bilayer before and after 62 min of incubation with Srf (3 µM).

(a)	(b)

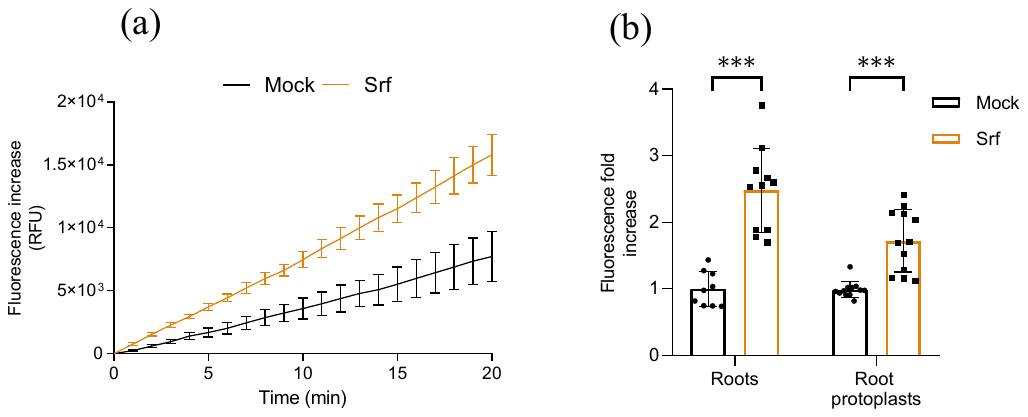

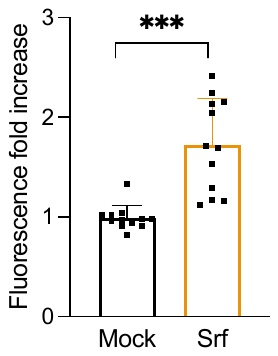

**Suppl Fig 15. (a)** Kinetic of [ROS]_intra_ production triggered by Srf in *Arabidopsis* Col-0 root protoplasts and measured with the fluorescent probe DFCH-DA. Graphs represent mean fluorescence increase observed ± SD (n=4). Experiment was repeated four times with similar results. **(b)** Fold increase in [ROS]_intra_ production in protoplasts (n=14 from four independent experiments) of *Arabidopsis* Col-0. Data represents mean values ± SD recorded at 5 min after the addition of 10 µM Srf compared to values obtained for mock-treated protoplasts. Asterisks indicate significant difference (****P*<0.001, two-tailed *t*-test).

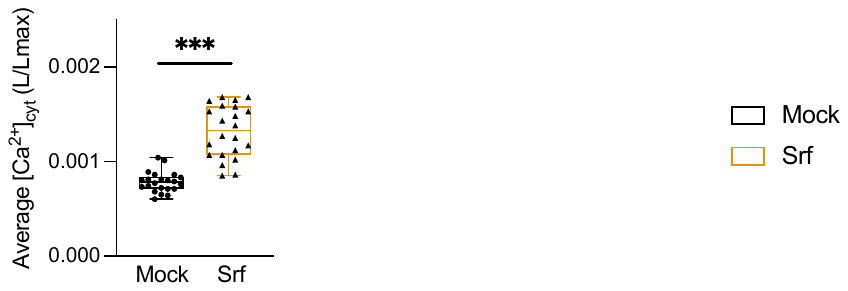

Suppl Fig 16. [Ca^2+^]_cyt_ response in Srf-treated (10 μM) root cell protoplasts (n=6) compared to mock controls in the *Arabidopsis* Col-0*^AEQ^* reporter line. Results are represented as luminescence counts per second relative to total luminescence counts remaining (L/Lmax). Values are the average of L/Lmax values from 1.5 to 4 min after treatment corresponding to the top of the peak. Mean ± SD of 14 replicates from four independent experiments. Asterisks indicate significant difference (****P*<0.001, two-tailed *t*-test).

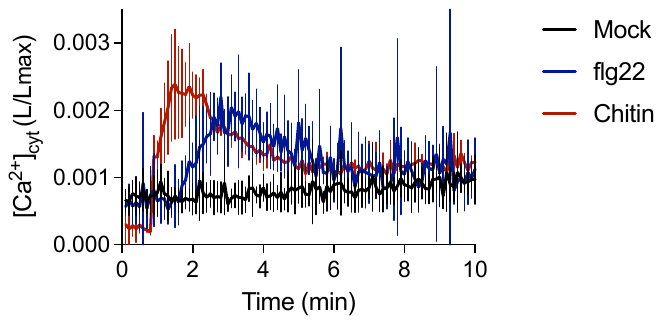

**Suppl Fig 17.** Kinetics of [Ca^2+^]_cyt_ production in *Arabidopsis* Col-0^AEQ^ root protoplasts treated with 1 µM Flg22 and 100 µg/mL chitin. Results are presented as luminescence counts per second relative to total luminescence counts remaining (L/Lmax) ± SD, n=8 from two independent experiments.

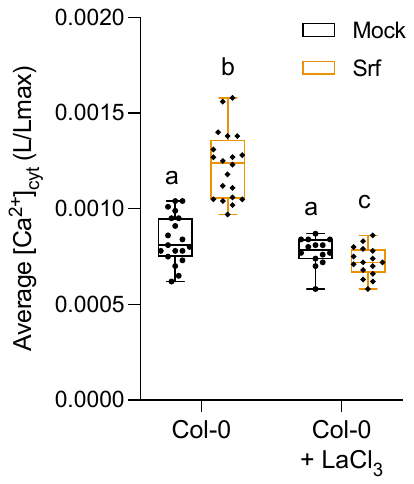

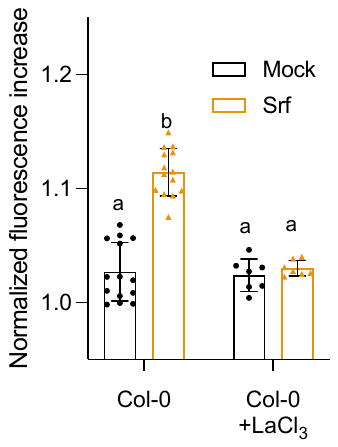

**Suppl Fig 18.** Effect of pre-treatment with the channel blocker LaCl_3_ (10 mM) on calcium response triggered by Srf (10 μM) in root protoplasts. Left, [Ca^2+^]_cyt_ increase measured in the Col-0*^AEQ^* reporter line. Results are represented as luminescence counts per second relative to total luminescence counts remaining (L/Lmax). Values are the average of L/Lmax values from 1.5 to 4 min after treatment corresponding to the top of the peak. Mean ± SD of 16 replicates from four independent experiments. Right, calcium influx measured via the Fluo-4 fluorescence probe in Col-0 protoplasts. Data are represented as mean normalized fluorescence increase (± SD) of 12 replicates from four independent experiments (for protoplasts without LaCl_3_ treatment) and of seven replicates from two independent experiments (for protoplasts with LaCl_3_ treatment). Letters represent statistically different groups at α = 0.05 (two-way ANOVA and Tukey’s multiple-comparison post-test).

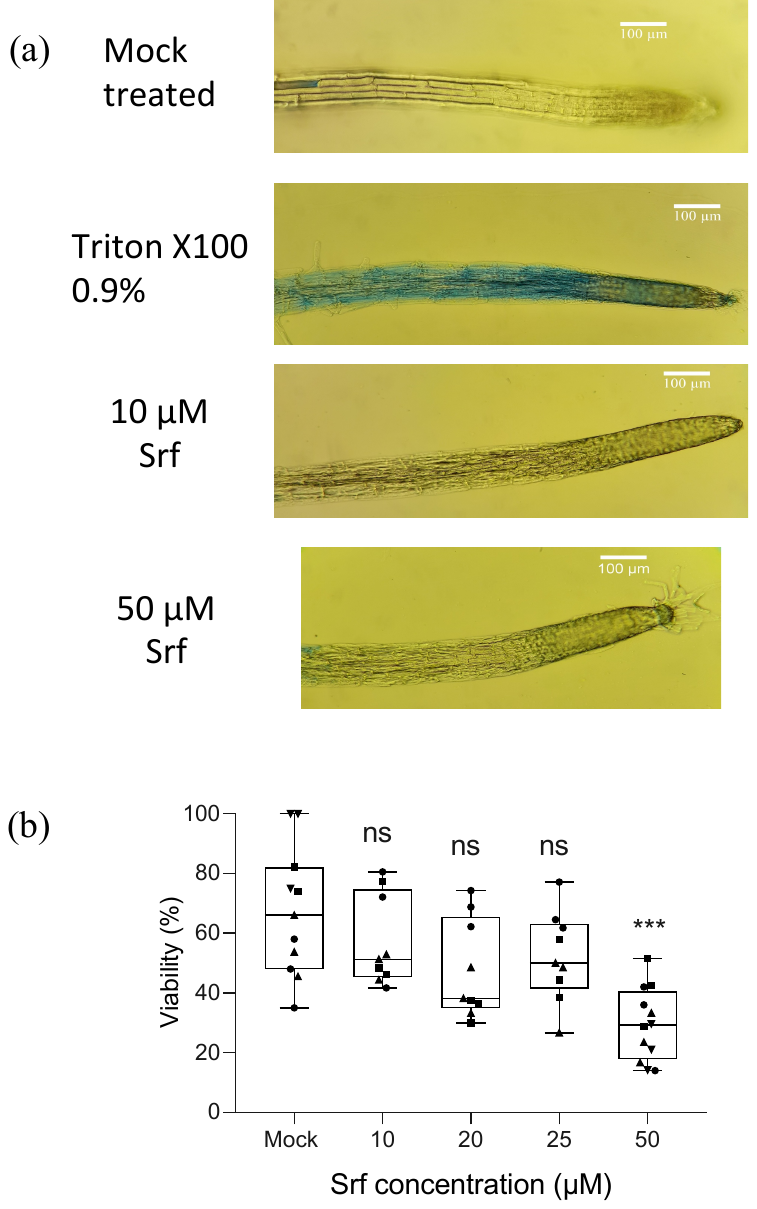

**Suppl Fig 19.** Viability of root protoplasts measured with fluorescein diacetate in the presence of increasing concentrations of Srf or 0.5% ethanol (Mock, negative control). Mean ± SD of 9 to 12 replicates (symbols on the graph) from three to four independent experiments (shown as different symbol shapes). Asterisks indicate statistically significant differences to the mock treatment (Brown-Forsythe and Welch ANOVA and Dunett’s T3 multiple comparisons test; ns, not significant; ****P* < 0.001).

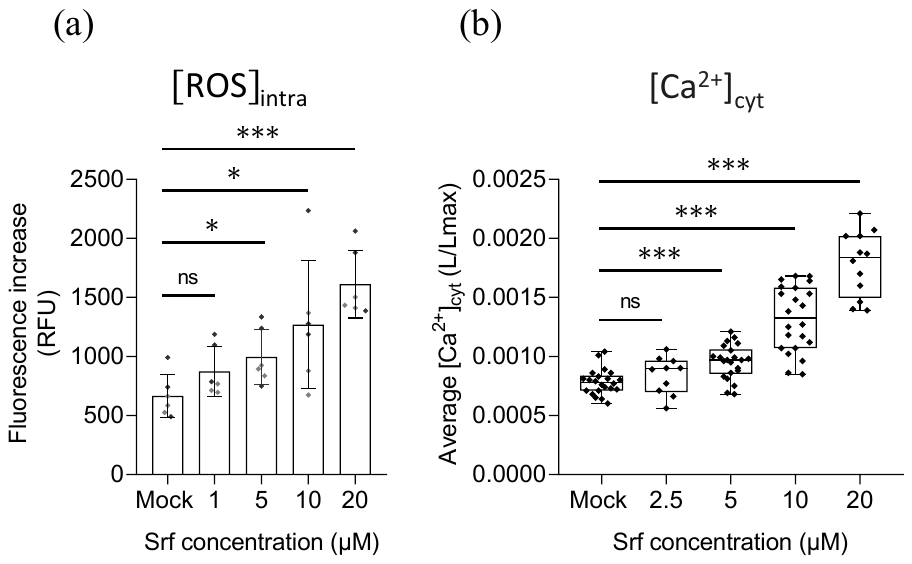

**Suppl Fig 20.** Dose-dependent [ROS]_intra_ induction by Srf in root protoplasts of *Arabidopsis*. Graph represents grouped data of two independent experiments (each n=3) presented as differently shaded grey values. Asterisks indicate statistically significant differences to the mock treatment (ns= no significant difference; **P*<0.05; ****P* < 0.001; (a) two-tailed *t*-test; (b) Welch and Brown-Forsythe ANOVA).

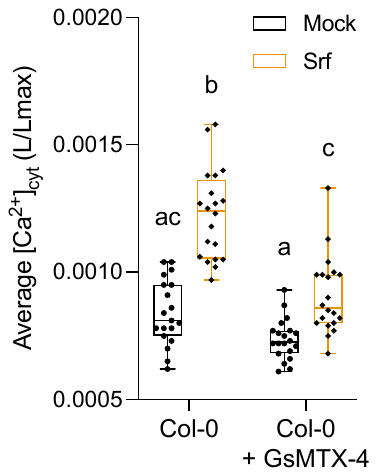

**Suppl Fig 21.** Effect of pre-treatment with the mechanosensitive channel blocker GsMTX-4 (10 min incubation, 7.5 µM) on [Ca^2+^]_cyt_ response in *Arabidopsis* Col-0*^AEQ^* upon Srf elicitation (10 µM). Data represent L/L_max_ values from 1.5 to 4 min after treatment corresponding to the top of the peak. Mean ± SD of at least 19 technical replicates from ten independent experiments. Letters represent statistically different groups at α = 0.05 (two-way ANOVA and Tukey’s multiple-comparison post-test).

**Supplementary table 1:** List of genes associated with plant immune responses used for the heatmap assembly in Fig 1. Log_2_ FC (Fold Change > 2, P<0.05) data for Srf are from the RNAseq analysis performed in this study while those reported for DEGs in response to flg22 (FLG) and chitin (Chi) are from published data^14^.

|  |  |  |  | **0.5 h** | **1 h** | **3 h** | **6 h** |
| --- | --- | --- | --- | --- | --- | --- | --- |
| **Calcium binding** | At1g76640 | Calcium-binding EF-hand family protein | SRF | -2.709561396 | 2.539243369 |  |  |
|  |  |  | FLG |  |  |  |  |
|  |  |  | CHI |  |  |  |  |
|  | At4g27280 | Calcium-binding EF-hand family protein | SRF |  |  |  | -3.323327514 |
|  |  |  | FLG |  | 2.028331985 |  |  |
|  |  |  | CHI | 2.621776197 |  |  |  |
|  | At5g39670 | Calcium-binding EF-hand family protein | SRF |  |  |  | -2.08716 |
|  |  |  | FLG | 4.079255898 | 3.080231186 | 2.421591421 | 3.468033038 |
|  |  |  | CHI | 5.355079351 |  | 2.050672768 |  |
|  | At3g47480 | Calcium-binding EF-hand family protein | SRF |  | -2.020426711 |  |  |
|  |  |  | FLG |  |  |  | 3.956896284 |
|  |  |  | CHI |  |  |  |  |
|  | At3g29000 | Calcium-binding EF-hand family protein | SRF |  |  |  | -3.141157853 |
|  |  |  | FLG | 3.486049655 | 3.949282548 | 2.433539561 | 3.172081877 |
|  |  |  | CHI | 5.845523475 |  |  |  |
|  | At1g73805 | Calmodulin binding protein-like | SRF |  |  |  |  |
|  |  |  | FLG | 3.069783141 |  | 2.239861705 | 3.387550993 |
|  |  |  | CHI | 4.397575173 |  |  |  |
|  | At2g41090 | Calcium-binding EF-hand family protein | SRF |  |  |  |  |
|  |  |  | FLG |  | 2.013554498 |  |  |
|  |  |  | CHI |  |  |  |  |
|  | At2g41100 | ATCAL4, TCH3, Calcium-binding EF hand family protein | SRF |  |  |  |  |
|  |  |  | FLG | 3.926611064 | 4.422389247 | 4.461846569 | 3.805786906 |
|  |  |  | CHI | 5.249806798 | 3.652430514 | 2.127493966 |  |
|  | At3g01830 | Calcium-binding EF-hand family protein | SRF |  |  |  |  |
|  |  |  | FLG | 4.937935778 | 3.966521775 | 2.591005708 | 2.581869329 |
|  |  |  | CHI | 5.822795741 |  |  |  |
|  | At1g76650 | CML38, calmodulin-like 38 | SRF |  |  |  |  |
|  |  |  | FLG |  | 3.459973174 | 2.29587099 | 3.215909601 |
|  |  |  | CHI | 3.610950882 | 2.839974679 |  |  |
|  | At4g20780 | CML42, calmodulin like 42 | SRF |  |  |  |  |
|  |  |  | FLG | 2.580067828 | 2.791439458 | 2.58073498 | 2.817497968 |
|  |  |  | CHI | 3.838220963 |  |  |  |
|  | At4g33050 | EDA39, calmodulin-binding family protein | SRF |  |  |  |  |
|  |  |  | FLG | 3.153435903 | 3.678456133 | 3.615381012 | 4.018451812 |
|  |  |  | CHI | 4.62844976 | 2.196147104 | 2.099137965 |  |
|  | At1g21550 | Calcium-binding EF-hand family protein | SRF |  |  |  |  |
|  |  |  | FLG |  |  |  | 2.148163074 |
|  |  |  | CHI | 2.025351244 |  |  |  |
|  | At3g51920 | ATCML9, CAM9, CML9, calmodulin 9 | SRF |  |  |  |  |
|  |  |  | FLG |  |  | 2.157218446 |  |
|  |  |  | CHI | 2.530785286 |  |  |  |
|  | At1g29025 | Calcium-binding EF-hand family protein | SRF |  |  |  |  |
|  |  |  | FLG |  | -2.133585656 | -2.654334669 | -4.453621149 |
|  |  |  | CHI |  |  |  |  |
|  | At1g29020 | Calcium-binding EF-hand family protein | SRF |  |  |  |  |
|  |  |  | FLG |  |  |  | -3.037667761 |
|  | At1g24620 | EF hand calcium-binding protein family | SRF |  |  |  |  |
|  |  |  | FLG |  |  |  | -3.796654813 |
|  | At1g05990 | RHS1, EF hand calcium-binding protein family | SRF |  |  |  |  |
|  |  |  | FLG |  |  |  | -4.214541416 |
|  | At3g59370 | Vacuolar calcium-binding protein-related | SRF |  |  |  |  |
|  |  |  | FLG |  |  |  | -3.823297994 |
|  | At2g24300 | Calmodulin-binding protein | SRF |  |  |  |  |
|  |  |  | FLG |  |  |  | -2.215335546 |
|  | At2g17890 | CPK16, calcium-dependent protein kinase 16 | SRF |  |  |  |  |
|  |  |  | FLG |  | -2.086250975 | -4.687855838 | -4.854535883 |
|  | At1g61950 | CPK19, calcium-dependent protein kinase 19 | SRF |  |  |  |  |
|  |  |  | FLG |  |  |  | -2.272654092 |
|  | At4g04710 | CPK22, calcium-dependent protein kinase 22 | SRF |  |  |  |  |
|  |  |  | FLG |  |  |  | -2.425083908 |
|  |  |  | CHI | 2.149769199 |  |  |  |
|  | At2g35890 | CPK25, calcium-dependent protein kinase 25 | SRF |  |  |  |  |
|  |  |  | FLG |  |  |  | -2.458046029 |
|  | At4g04700 | CPK27, calcium-dependent protein kinase 27 | SRF |  |  |  |  |
|  |  |  | FLG |  | 2.209407288 |  |  |
|  |  |  | CHI | 2.745107415 |  |  |  |
|  | At5g66210 | CPK28, calcium-dependent protein kinase 28 | SRF |  |  |  |  |
|  |  |  | FLG | 2.578481147 |  |  |  |
|  |  |  | CHI | 3.636901982 |  |  |  |
|  | At5g42380 | CML37, CML39, calmodulin like 37 | SRF |  |  |  | -3.5884293 |
|  |  |  | CHI | 6.186147103 | 2.987318816 |  |  |
|  | AT5G57010 | calmodulin-binding family protein | SRF |  |  |  | -2.059418183 |
|  |  |  | CHI | 2.083817721 |  |  |  |
|  | At2g26190 | calmodulin-binding family protein | SRF |  |  |  |  |
|  |  |  | CHI | 2.480707971 |  |  |  |
|  | At3g25600 | Calcium-binding EF-hand family protein | SRF |  |  |  |  |
|  |  |  | CHI | 2.214664142 |  |  |  |
|  | At3g57530 | ATCPK32, CDPK32, CPK32, calcium-dependent protein kinase 32 | SRF |  |  |  |  |
|  |  |  | CHI | 2.408235454 |  |  |  |
| ROS | AT1G14550 | Peroxidase superfamily protein (extracellular region) | SRF | 3.002402118 | 5.612086065 | 4.723695277 | 6.710333507 |
|  | EXTR |  | FLG | 7.499396788 | 8.111306245 | 8.319699362 | 8.861195573 |
|  |  |  | chi | 8.70282543 | 5.97550331 | 3.449174822 | 3.466223449 |
|  | AT5G05340 | Peroxidase superfamily protein | SRF |  |  |  | 6.200332294 |
|  |  |  | FLG |  |  | 5.515805668 | 8.508524809 |
|  | AT1G49570 | Peroxidase superfamily protein | SRF |  |  | 2.046154573 | 3.546594762 |
|  |  |  | FLG |  |  |  | 3.269697061 |
|  | AT5G64120 | Peroxidase superfamily protein | SRF |  |  |  | 3.129211457 |
|  |  |  | FLG |  | 1.662221072 | 2.733938568 | 3.097878429 |
|  | AT1G14540 | Peroxidase superfamily protein (extracellular region) | SRF | 3.307228769 | 2.483080377 | 4.056903911 | 2.807897066 |
|  | CELL WALL |  | FLG | 5.353558128 | 5.868604702 | 5.788059579 | 5.571581885 |
|  |  |  | chi | 7.146150443 | 4.510168027 | 2.248612035 | 2.321544033 |
|  | AT5G19890 | Peroxidase superfamily protein | SRF |  |  |  | 2.541034038 |
|  |  |  | FLG |  |  | 2.255415645 | 3.505073627 |
|  | AT3G03670 | Peroxidase superfamily protein | SRF |  |  | 2.034061669 | 2.083661278 |
|  |  |  | FLG |  | 2.922751192 | 4.936070939 | 7.024892739 |
|  |  |  | chi | 2.96855382 | 3.975609514 | 3.551150699 |  |
|  | AT5G64110 | Peroxidase superfamily protein | SRF |  |  | 2.354189092 |  |
|  |  |  | FLG |  |  | 2.492034844 | 2.813359013 |
|  | AT2G41480 | Peroxidase superfamily protein | SRF |  |  |  |  |
|  |  |  | FLG |  |  | -2.013040238 |  |
|  | AT5G39580 | Peroxidase superfamily protein | FLG | 5.513418715 | 5.794345215 | 6.895428137 | 7.009066439 |
|  |  |  | chi | 6.404309798 | 4.979448157 | 3.501955947 | 2.605441767 |
|  | AT3G49110 | ATPCA, ATPRX33, PRX33, PRXCA, peroxidase CA | chi | 3.644207014 | 4.563256138 | 2.565611546 |  |
|  |  |  | FLG |  | 4.393840729 | 5.296369639 | 7.393304925 |
|  | AT5G19880 | Peroxidase superfamily protein | flg |  | 4.166038549 | 5.447068954 | 5.203287776 |
|  |  |  | chi | 2.230593394 | 4.717422393 | 3.22814285 | 4.998498104 |
|  | AT4g11290 | Peroxidase superfamily protein | flg |  |  |  | 2.754164942 |
|  | AT4g08780 | Peroxidase superfamily protein | FLG |  |  | 3.745741048 | 5.809665998 |
|  |  |  | chi |  | 2.01941591 |  |  |
|  | AT5g15180 | Peroxidase superfamily protein | FLG |  | -2.488657182 | -3.925955351 | -3.825011453 |
|  | AT1g30870 | Peroxidase superfamily protein | FLG |  |  |  | -2.863452503 |
|  | AT2g39040 | Peroxidase superfamily protein | FLG |  |  | -2.22781561 | -5.443705565 |
|  | AT4g08770 | Peroxidase superfamily protein | FLG |  |  | 2.514109825 | 3.419325491 |
|  |  |  | chi |  |  |  |  |
|  | AT3g49960 | Peroxidase superfamily protein | FLG |  |  |  | -4.740568272 |
|  | AT1g34510 | Peroxidase superfamily protein | FLG |  |  |  | -3.268622544 |
|  | AT1g71695 | Peroxidase superfamily protein | FLG |  |  |  | 4.001744665 |
|  | AT2g43480 | Peroxidase superfamily protein | FLG |  |  |  | -4.792135631 |
|  | AT4g37520 | Peroxidase superfamily protein | FLG |  |  |  | 3.465365591 |
|  | AT2g18980 | Peroxidase superfamily protein | FLG |  |  |  | -2.109458135 |
|  | AT4g16270 | Peroxidase superfamily protein | FLG |  | -2.396516788 |  | -3.68935469 |
|  | AT5G07390 | ATRBOHA, RBOHA, respiratory burst oxidase homolog A | SRF |  |  |  |  |
|  |  |  | FLG |  |  | 2.340460316 | 2.54166697 |
|  |  |  | chi | 2.354323009 |  |  |  |
|  | AT5G47910 | ATRBOHD, RBOHD, respiratory burst oxidase homologue D | SRF |  |  |  |  |
|  |  |  | FLG |  |  |  | 2.014832148 |
|  |  |  | chi | 2.890414113 |  |  |  |
| **MAPK** | At1g18350 | ATMKK7, BUD1, MKK7, MKK7, MAP kinase kinase 7 | SRF |  |  |  |  |
|  |  |  | FLG |  |  | 3.752727738 | 5.394811887 |
|  | At1g73500 | ATMKK9, MKK9, MAP kinase kinase 9 | SRF |  |  |  |  |
|  |  |  | FLG |  | 3.599656239 | 2.572877933 | 3.250302553 |
|  |  |  | CHI |  | 2.798543399 |  |  |
|  | At5g55090 | MAPKKK15, mitogen-activated protein kinase kinase kinase 15 | SRF |  |  |  |  |
|  |  |  | FLG | 3.230208634 | 2.749673751 | 2.471953929 | 1.898647311 |
|  |  |  | CHI | 2.713282065 |  |  |  |
|  | At5g67080 | MAPKKK19, mitogen-activated protein kinase kinase kinase 19 | SRF |  |  |  |  |
|  |  |  | FLG |  | 3.047285554 | 3.950138774 | 4.979043047 |
|  |  |  | CHI | 2.000291971 | 2.299997703 |  |  |
|  | At3g50310 | MAPKKK20, mitogen-activated protein kinase kinase kinase 20 | SRF |  |  |  |  |
|  |  |  | FLG |  |  | 2.413266841 | 4.094003602 |
|  | At1g01560 | ATMPK11, MPK11, MAP kinase 11 | SRF |  |  |  |  |
|  |  |  | FLG | 2.348282199 | 2.681554951 | 3.283135671 | 3.979556791 |
|  |  |  | CHI | 5.404437612 | 3.070886837 | 3.170447656 | 2.305258286 |
|  | At2g46070 | ATMPK12, MAPK12, MPK12, mitogen-activated protein kinase 12 | SRF |  |  |  |  |
|  |  |  | FLG |  |  |  | 2.004977991 |
|  | At3g45640 | ATMAPK3, ATMPK3, MPK3, mitogen-activated protein kinase 3 | SRF |  |  |  |  |
|  |  |  | flg |  |  |  |  |
|  |  |  | CHI | 2.141231255 |  |  |  |
|  | At1g07150 | MAPKKK13, mitogen-activated protein kinase kinase kinase 13 | srf |  |  |  |  |
|  |  |  | FLG |  |  |  | 2.256901913 |
| **PR** | AT1g75830 | PDF1.1, LCR67, low-molecular-weight cysteine-rich 67 | SRF |  |  |  |  |
|  |  |  | FLG |  | 4.610594454 | 5.125465599 | 6.605854626 |
|  | AT5g44420 | PDF1.2, LCR77,PDF1.2A, plant defensin 1.2 | SRF |  |  |  |  |
|  |  |  | FLG |  |  |  | 6.546291931 |
|  | AT2g26020 | PDF1.2b, plant defensin 1.2b | SRF |  |  |  |  |
|  |  |  | FLG |  |  | 3.673731539 | 5.162284645 |
|  | AT1g19610 | LCR78, PDF1.4, Arabidopsis defensin-like protein | SRF |  |  |  |  |
|  |  |  | FLG |  |  | 2.374293493 | 3.094867892 |
|  |  |  | chi |  |  | 2.361262476 |  |
|  | AT2g02120 | LCR70, PDF2.1, Scorpion toxin-like knottin superfamily protein | SRF | -2.499475259 |  |  |  |
|  |  |  | FLG |  |  |  | -2.775078036 |
|  | AT5g26130 | CAP (Cysteine-rich secretory proteins, Antigen 5, and Pathogenesis-related 1 protein) superfamily protein | SRF |  |  |  |  |
|  |  |  | FLG |  |  |  | 4.242856251 |
|  | AT4g33710 | CAP (Cysteine-rich secretory proteins, Antigen 5, and Pathogenesis-related 1 protein) superfamily protein | SRF |  |  |  |  |
|  |  |  | FLG |  |  |  | 4.797242254 |
|  |  |  | chi |  |  | 4.220003068 | 4.523308265 |
|  | AT4g25790 | CAP (Cysteine-rich secretory proteins, Antigen 5, and Pathogenesis-related 1 protein) superfamily protein | SRF |  |  |  |  |
|  |  |  | FLG |  |  |  | -4.258810254 |
|  | AT5g57625 | CAP (Cysteine-rich secretory proteins, Antigen 5, and Pathogenesis-related 1 protein) superfamily protein | SRF |  |  |  |  |
|  |  |  | FLG |  |  |  | -2.629055212 |
|  | AT4g30320 | CAP (Cysteine-rich secretory proteins, Antigen 5, and Pathogenesis-related 1 protein) superfamily protein | SRF |  |  |  |  |
|  |  |  | FLG |  |  |  | -2.527777314 |
|  | AT4g31470 | CAP (Cysteine-rich secretory proteins, Antigen 5, and Pathogenesis-related 1 protein) superfamily protein | SRF |  |  |  |  |
|  |  |  | FLG |  |  |  | -4.044894901 |
|  | AT5g66590 | CAP (Cysteine-rich secretory proteins, Antigen 5, and Pathogenesis-related 1 protein) superfamily protein | SRF |  |  |  |  |
|  |  |  | FLG |  |  | -2.716612805 | -5.291280318 |
|  | AT2g19980 | CAP (Cysteine-rich secretory proteins, Antigen 5, and Pathogenesis-related 1 protein) superfamily protein | SRF |  |  |  |  |
|  |  |  | chi | -4.59117346 |  |  |  |
|  | AT3g04720 | HEL, PR-4, PR4, pathogenesis-related 4 | SRF |  |  |  |  |
|  |  |  | FLG |  |  |  | 2.713256135 |
|  | AT1g73620 | Pathogenesis-related thaumatin superfamily protein | SRF |  |  |  |  |
|  |  |  | FLG |  |  |  | -3.543760799 |
|  | AT4g38660 | Pathogenesis-related thaumatin superfamily protein | SRF |  |  |  |  |
|  |  |  | FLG |  |  |  | -2.054527891 |
|  | AT2g28790 | Pathogenesis-related thaumatin superfamily protein | SRF |  |  |  |  |
|  |  |  | FLG |  |  |  | -2.333400225 |
|  | AT2g19990 | PR-1-LIKE, pathogenesis-related protein-1-like | SRF |  |  |  |  |
|  |  |  | FLG |  |  |  | -3.973094345 |
|  | AT3g12500 | ATHCHIB, B-CHI, CHI-B, HCHIB, PR-3, PR3, basic chitinase | SRF |  |  |  |  |
|  |  |  | FLG |  |  |  | 2.249474035 |
|  | AT5g24090 | ATCHIA, CHIA, chitinase A | SRF |  |  |  |  |
|  |  |  | FLG |  |  |  | 2.077478016 |
|  | AT3g54420 | ATCHITIV, ATEP3, CHIV, EP3, homolog of carrot EP3-3 chitinase | SRF |  |  |  | 2.13386159 |
|  |  |  | FLG | 2.079851905 | 2.289309101 |  |  |
|  |  |  | chi | 2.906198149 |  |  |  |
|  | AT4g01700 | Chitinase family protein | SRF |  |  |  |  |
|  |  |  | FLG |  | 2.973675974 | 3.115226327 | 2.363776878 |
|  |  |  | chi | 2.727723618 | 2.713399178 |  |  |
|  | AT1g56680 | Chitinase family protein | SRF |  |  |  |  |
|  |  |  | FLG |  |  |  | -2.583965659 |
|  | AT2g43590 | Chitinase family protein | SRF |  |  | 2.229177716 | 2.106874527 |
|  |  |  | FLG |  |  |  | 2.249936719 |
|  | AT1g02360 | Chitinase family protein | SRF |  |  |  |  |
|  |  |  | FLG |  | 2.292684172 | 2.228147909 |  |
|  |  |  | chi | 2.641940641 | 2.149419752 |  |  |
|  | AT2g43620 | Chitinase family protein | SRF |  |  |  |  |
|  |  |  | FLG | 5.194373127 | 7.068849608 | 7.776061838 | 5.915189329 |
|  |  |  | chi | 3.844437333 | 4.447878651 | 3.89581102 | 2.648728138 |
|  | AT2g43580 | Chitinase family protein | SRF |  |  |  |  |
|  |  |  | FLG |  |  |  | 2.575243624 |
|  |  |  | chi | 2.145213514 | 2.053854712 |  |  |
|  | AT3g47540 | Chitinase family protein | chi | 2.187025186 |  |  |  |
| **Receptor-like kinases** | At1g65790 | ARK1, RK1, receptor kinase 1 | SRF |  |  |  |  |
|  |  |  | FLG |  |  | 7.037175041 | 4.34974188 |
|  |  |  |  | 6.141874138 |  | 6.734360808 | 4.812048175 |
|  | At1g65800 | ARK2, RK2, receptor kinase 2 | SRF |  |  |  |  |
|  |  |  | chi | 3.594359127 |  |  |  |
|  | At4g21380 | ARK3, RK3, receptor kinase 3 | SRF |  |  |  |  |
|  |  |  | FLG | 2.845850445 | 4.340555281 | 5.883664016 | 6.040672878 |
|  |  |  | chi | 4.994693718 | 5.659028003 | 4.702779445 | 4.721433107 |
|  | At5g48400 | ATGLR1.2, GLR1.2, Glutamate receptor family protein | SRF |  |  |  |  |
|  |  |  | FLG |  |  |  | 3.412910967 |
|  |  |  | chi | 3.963965974 | 3.079804701 | 2.516142813 | 2.351960483 |
|  | At5g48410 | ATGLR1.3, GLR1.3, glutamate receptor 1.3 | SRF |  |  |  |  |
|  |  |  | FLG |  |  |  | 2.660921 |
|  |  |  | chi | 2.314472886 |  |  |  |
|  | At5g11210 | ATGLR2.5, GLR2.5, glutamate receptor 2.5 | SRF |  |  |  |  |
|  |  |  | FLG | 3.93915936 | 2.382394239 | 3.348402552 | 3.757952025 |
|  |  |  | chi | 4.495972384 |  |  | 2.055210527 |
|  | At2g29110 | ATGLR2.8, GLR2.8, GLR2.8, glutamate receptor 2.8 | SRF |  |  |  |  |
|  |  |  | FLG | 2.710447519 | 2.578679948 | 2.714963577 | 2.573733339 |
|  |  |  | chi | 5.743486518 | 2.570028968 |  |  |
|  | At5g11180 | ATGLR2.6, GLR2.6, glutamate receptor 2.6 | SRF |  |  |  |  |
|  |  |  | chi | 2.81845743 |  |  |  |
|  | At2g29120 | ATGLR2.7, GLR2.7, GLR2.7, glutamate receptor 2.7 | SRF |  |  |  |  |
|  |  |  | chi | 2.793172418 |  | 2.137894262 |  |
|  | At2g29100 | ATGLR2.9, GLR2.9, GLR2.9, glutamate receptor 2.9 | SRF |  |  |  |  |
|  |  |  | FLG | 3.718977136 | 3.118631941 | 3.587129824 | 3.91645606 |
|  |  |  | chi | 5.807821695 | 3.717777547 | 2.673688884 | 2.799836252 |
|  | At3g59700 | ATHLECRK, HLECRK, LECRK1, lectin-receptor kinase | SRF |  |  |  |  |
|  |  |  | FLG |  |  |  | 2.303704207 |
|  |  |  | chi | 2.091837411 |  |  |  |
|  | At4g23190 | AT-RLK3, CRK11, cysteine-rich RLK (RECEPTOR-like protein kinase) 11 | SRF |  |  |  |  |
|  |  |  | FLG | 3.051427512 | 2.220036137 | 2.442321754 | 2.421928133 |
|  |  |  | chi | 3.6949108 |  |  |  |
|  | At5g58940 | CRCK1, calmodulin-binding receptor-like cytoplasmic kinase 1 | FLG |  |  |  |  |
|  |  |  | chi | 2.040420344 |  |  |  |
|  | At4g00330 | CRCK2, calmodulin-binding receptor-like cytoplasmic kinase 2 | chi | 2.042141021 |  |  |  |
|  | At4g23180 | CRK10, RLK4, cysteine-rich RLK (RECEPTOR-like protein kinase) 10 | SRF |  |  |  |  |
|  |  |  | FLG | 3.969912773 | 2.678073859 | 2.961898467 | 2.192467296 |
|  |  |  | chi | 3.83588765 | 2.862944852 | 2.087998038 |  |
|  | At4g23220 | CRK14, cysteine-rich RLK (RECEPTOR-like protein kinase) 14 | SRF |  |  |  |  |
|  |  |  | FLG | 3.975306353 | 4.553911723 | 4.584776985 | 4.148590386 |
|  |  |  | chi | 4.989028204 | 5.551430487 | 2.757692524 |  |
|  | At4g23230 | CRK15, cysteine-rich RLK (RECEPTOR-like protein kinase) 15 | SRF |  |  |  |  |
|  |  |  | FLG |  | 3.912289173 |  |  |
|  |  |  | chi |  | 3.904599342 |  |  |
|  | At4g23250 | CRK17, DUF26-21, EMB1290, RKC1, kinases;protein kinases | SRF |  |  |  |  |
|  |  |  | FLG | 3.45791659 | 3.926957648 | 3.510820778 | 3.030563871 |
|  |  |  | chi | 4.979207333 | 4.519206283 |  |  |
|  | At4g23260 | CRK18, cysteine-rich RLK (RECEPTOR-like protein kinase) 18 | SRF |  |  |  |  |
|  |  |  | FLG |  | 2.451005789 | 3.572035897 | 3.290313686 |
|  |  |  | chi |  | 2.119640358 |  |  |
|  | At4g23270 | CRK19, cysteine-rich RLK (RECEPTOR-like protein kinase) 19 | chi | 2.076480588 |  |  |  |
|  | At4g23280 | CRK20, cysteine-rich RLK (RECEPTOR-like protein kinase) 20 | SRF |  |  |  |  |
|  |  |  | FLG | 3.172431318 | 5.473728457 | 6.137296133 | 5.006810988 |
|  |  |  | chi | 6.015166079 | 5.721204844 | 4.532239307 | 3.208017268 |
|  | At5g01550 | LECRKA4.2, lectin receptor kinase a4.1 | SRF |  | 2.078076237 |  |  |
|  |  |  | FLG |  |  |  |  |
|  |  |  | chi | 2.789862378 |  |  |  |
|  | At4g23300 | CRK22, cysteine-rich RLK (RECEPTOR-like protein kinase) 22 | SRF |  | -4.47190577 | -4.861023043 | 4.982655416 |
|  |  |  | FLG |  |  |  |  |
|  | At4g23320 | CRK24, cysteine-rich RLK (RECEPTOR-like protein kinase) 24 | SRF |  |  |  |  |
|  |  |  | FLG |  |  |  | 5.341150864 |
|  |  |  | chi | 4.228130831 |  |  |  |
|  | At4g38830 | CRK26, cysteine-rich RLK (RECEPTOR-like protein kinase) 26 | chi | 2.135646764 |  |  |  |
|  | At4g21400 | CRK28, cysteine-rich RLK (RECEPTOR-like protein kinase) 28 | SRF |  |  |  |  |
|  |  |  | FLG |  |  |  | 2.503823868 |
|  | At4g11470 | CRK31, cysteine-rich RLK (RECEPTOR-like protein kinase) 31 | SRF |  |  |  |  |
|  |  |  | FLG | 3.484559971 | 3.454108643 | 3.671722574 | 3.672487247 |
|  |  |  | chi | 5.791438179 | 3.353937428 |  |  |
|  | At4g11480 | CRK32, cysteine-rich RLK (RECEPTOR-like protein kinase) 32 | SRF |  |  |  |  |
|  |  |  | FLG |  |  | 2.559450232 |  |
|  |  |  | chi | 4.184371596 | 3.347733599 |  |  |
|  | At4g04540 | CRK39, cysteine-rich RLK (RECEPTOR-like protein kinase) 39 | SRF |  |  |  |  |
|  |  |  | FLG |  |  | 7.50941361 | 10.58660697 |
|  |  |  | chi |  |  | 4.674851761 | 5.580542508 |
|  | At4g04570 | CRK40, cysteine-rich RLK (RECEPTOR-like protein kinase) 40 | SRF |  |  |  |  |
|  |  |  | FLG |  |  | 2.658833522 | 2.927020249 |
|  |  |  | chi | 4.247448622 | 2.541422496 | 2.181188593 | 2.124443684 |
|  | At4g00970 | CRK41, cysteine-rich RLK (RECEPTOR-like protein kinase) 41 | SRF |  |  |  |  |
|  |  |  | FLG | 5.255124369 | 4.175324317 | 4.582952087 | 3.683174405 |
|  |  |  | chi | 6.914747258 | 5.687247816 | 2.782495568 |  |
|  | At4g23130 | CRK5, RLK6, cysteine-rich RLK (RECEPTOR-like protein kinase) 5 | SRF |  |  |  |  |
|  |  |  | FLG |  | 2.186805947 |  |  |
|  | At4g23140 | CRK6, cysteine-rich RLK (RECEPTOR-like protein kinase) 6 | SRF |  |  |  |  |
|  |  |  | FLG |  |  | 3.14608972 |  |
|  | At4g23170 | CRK9, EP1, receptor-like protein kinase-related family protein | SRF |  |  |  |  |
|  |  |  | FLG | 3.904997973 | 3.46768288 | 3.647216077 | 2.338574873 |
|  |  |  | chi | 4.5638763 | 3.455254545 | 2.791777269 |  |
|  | At5g01540 | LECRKA4.1, lectin receptor kinase a4.1 | SRF |  |  |  |  |
|  |  |  | FLG | 3.789411728 | 4.66052121 | 3.999668194 | 2.699611622 |
|  |  |  | chi | 4.640136182 | 2.007144258 |  |  |
|  | At5g01560 | LECRKA4.3, lectin receptor kinase a4.3 | SRF |  |  |  |  |
|  |  |  | FLG |  | 2.432734424 | 2.114612684 | 2.054293499 |
|  |  |  | chi | 2.042856939 |  |  |  |
|  | At4g20940 | Leucine-rich receptor-like protein kinase family protein | SRF |  |  |  |  |
|  |  |  | FLG |  |  |  | -2.251724484 |
|  |  |  | chi |  |  |  |  |
|  | At2g24130 | Leucine-rich receptor-like protein kinase family protein | SRF |  |  |  |  |
|  |  |  | FLG |  | 2.665702511 | 2.739210192 | 3.186400869 |
|  |  |  | chi |  | 2.250887961 |  |  |
|  | At3g28040 | Leucine-rich receptor-like protein kinase family protein | SRF |  |  |  |  |
|  |  |  | FLG |  |  |  | -3.608213772 |
|  |  |  | chi |  |  |  |  |
|  | At5g01890 | Leucine-rich receptor-like protein kinase family protein | SRF |  |  |  |  |
|  |  |  | FLG |  |  |  | -2.072978506 |
|  |  |  | chi |  |  |  |  |
|  | At5g46330 | FLS2, Leucine-rich receptor-like protein kinase family protein | SRF |  |  |  |  |
|  |  |  | FLG | 3.2858675 | 3.086817047 | 2.128688729 |  |
|  |  |  | chi | 2.003986919 |  |  |  |
|  | At2g19190 | FRK1, FLG22-induced receptor-like kinase 1 | SRF |  |  |  |  |
|  |  |  | FLG | 3.428550803 | 5.007094376 | 5.594355957 | 4.869353893 |
|  |  |  | chi |  | 3.003984424 |  |  |
|  | At5g48540 | receptor-like protein kinase-related family protein | SRF |  | 2.715678974 |  |  |
|  |  |  | FLG | 3.323522021 | 3.053157796 | 2.937792091 | 3.358995208 |
|  |  |  | chi | 3.386848396 | 2.267031113 |  |  |
|  | At1g63590 | Receptor-like protein kinase-related family protein | SRF |  |  | 2.546491844 |  |
|  |  |  | FLG |  | 2.17379688 | 3.691296978 | 4.345718177 |
|  |  |  | chi | 4.73770954 | 3.136344559 | 3.373780916 | 3.482461923 |
|  | At1g63570 | Receptor-like protein kinase-related family protein | SRF |  |  |  |  |
|  |  |  | FLG |  |  | 3.160494942 |  |
|  |  |  | chi | 3.870699033 | 3.801515015 | 4.412875322 |  |
|  | At5g38990 | Malectin/receptor-like protein kinase family protein | SRF |  |  | -3.969535972 |  |
|  |  |  | FLG | 2.071547437 | 2.325086498 |  |  |
|  | At1g63580 | Receptor-like protein kinase-related family protein | SRF |  |  |  | 2.054910564 |
|  |  |  | chi | 2.554517139 | 2.20760683 |  |  |
|  | At1g63560 | Receptor-like protein kinase-related family protein | SRF |  |  |  |  |
|  |  |  | FLG |  | 3.117599033 | 4.440902733 | 5.423295624 |
|  |  |  | chi | 2.965952987 | 6.614979302 | 5.001553156 | 5.00087594 |
|  | At1g63600 | Receptor-like protein kinase-related family protein | SRF |  |  |  |  |
|  |  |  | FLG |  |  | -3.945327016 | -6.013733779 |
|  |  |  | chi |  |  |  |  |
|  | At4g11521 | Receptor-like protein kinase-related family protein | SRF |  |  |  |  |
|  |  |  | FLG | 4.996571566 | 4.52353732 | 5.539894239 |  |
|  |  |  | chi | 6.708863419 | 3.53338467 | 3.85548764 |  |
|  | At5g60900 | RLK1, receptor-like protein kinase 1 | SRF |  |  |  |  |
|  |  |  | FLG |  |  | 3.134549646 | 2.782505965 |
|  |  |  | chi | 2.808241667 | 2.107577363 | 2.621998882 | 2.751524236 |
|  | At2g32140 | transmembrane receptors | SRF |  |  |  | -2.176622857 |
|  |  |  | FLG | 2.056826366 |  |  | 2.638448637 |
|  |  |  | chi | 3.497315853 |  |  |  |
|  | At3g05360 | AtRLP30, RLP30, receptor like protein 30 | SRF |  |  |  |  |
|  |  |  | FLG |  |  |  | 2.138137042 |
| **Cell wall** | AT1g67980 | CCOAMT, caffeoyl-CoA 3-O-methyltransferase | SRF |  |  |  |  |
|  |  |  | FLG | 2.026184727 | 3.580489967 | 5.82321927 | 6.958469021 |
|  |  |  | chi | 3.831938079 | 3.161275424 | 4.51098615 | 3.937110502 |
|  | AT1G80820 | ATCCR2, CCR2, cinnamoyl coa reductase | FLG |  |  |  | 2.497667669 |
|  |  |  | chi | 3.930437065 |  |  |  |
|  | AT4G36220 | CYP84A1, FAH1, ferulic acid 5-hydroxylase 1 | FLG |  |  | -3.618567945 | -3.660241069 |
|  | AT5g36870 | ATGSL09, atgsl9, gsl09, GSL09, glucan synthase-like 9 | FLG |  |  |  | -2.836790173 |
|  |  |  | chi |  |  |  | -2.080800792 |

**Supplementary Table 2**: List of primers used for qRT-PCR

| **Gene** | **Forward primer (5‘-3’)** | **Reverse primer (5’-3’)** |
| --- | --- | --- |
| *CML37* (At5g42380) | ACGAGCAGTAATAGTAGCGGAAGCA | CGCCTAAGAGACTAACGCAGCTTT |
| *CYP71A12* (At2g30750) | GATTATCACCTCGGTTCCT | CCACTAATACTTCCCAGATTA |
| *ERF6* (At4g17490) | GAAAACCGCCGTTGAAGATC | CGGTTGCGAATTGAATCCA |
| *MYB51* (At1G18570) | CTACAAGTGTTTCCGTTGACTCTGAA | ACGAAATTATCGCAGTACATTAGAGGA |
| *UBQ5*  (At3G62250) | TCTCCGTGGTGGTGCTAAG | GAACCTTTCCAGATCCATCG |

**Supplementary Table 3**: Relative amounts of the deuterated Srf C15 variant isotopes produced, the molecular formula of their fatty acid chain and peptide cycle.

|  | **FA non deut +**  **1 Leu deut** | **FA deut +**  **4 Leu non-deut.** | **FA not deut +**  **2 Leu deut** | **FA deut +**  **1 Leu deut** | **FA not deut +**  **3 Leu deut** | **FA deut +**  **2 Leu deut** | **FA not deut +**  **Leu not deut** |
| --- | --- | --- | --- | --- | --- | --- | --- |
| **Relative amount** | 0.261 | 0.100 | 0.260 | 0.077 | 0.222 | 0.047 | 0.034 |
| **FA** | C_12_H_25_ | C_12_H_16_D_9_ | C_12_H_25_ | C_12_H_16_D_9_ | C_12_H_25_ | C_12_H_16_D_9_ | C_12_H_25_ |
| **Cyclic peptide** | C_41_H_59_N_7_O_13_D_9_ | C_41_H_68_N_7_O_13_ | C_41_H_50_N_7_O_13_D_18_ | C_41_H_59_N_7_O_13_D_9_ | C_41_H_41_N_7_O_13_D_27_ | C_41_H_50_N_7_O_13_D_18_ | C_41_H_68_N_7_O_13_ |

**Supplementary table 4:** Structural parameters used to fit the Neutron reflectivity spectra relative to the **(a)** PLPC-Sito-GluCer bilayer and **(b)** PLPC-Sito bilayer before (left) and after (right) Srf addition. A contemporary fitting of data collected from the membranes in H_2_O and D_2_O has been performed. SLD: Scattering length density, FA: fatty acid chain.

**a**

|  | **PLPC-Sito-GluCer bilayer** | | | **PLPC-Sito-GluCer bilayer + Srf** | | |
| --- | --- | --- | --- | --- | --- | --- |
|  | Thickness  (±1Å) | SLD  (±0.05*10^-6^Å^-2^) | Solvent penetration  (±5%vol) | Thickness  (±1Å) | SLD  (±0.05*10^-6^Å^-2^) | Solvent penetration  (±5%vol) |
| **Heads in** | 6 | 1.93 | 25 | 6 | 1.93 | 25 |
| **Chains in** | 14 | -0.28 | 10 | 14 | -0.28 | 10 |
| **Chains out** | 14 | -0.41 | 10 | 10 | -0.38 | 20 |
| **Heads out** | 6 | 1.98 | 25 | 6 | 2 | 30 |

**b**

|  | | **PLPC-Sito bilayer** | | | | | **PLPC-Sito bilayer + Srf** | | | | |
| --- | --- | --- | --- | --- | --- | --- | --- | --- | --- | --- | --- |
|  | | Thickness  (±1Å) | | SLD  (±0.05*10^-6^Å^-2^) | | Solvent penetration  (±5%vol) | Thickness  (±1Å) | | SLD  (±0.05*10^-6^Å^-2^) | | Solvent penetration  (±5%vol) |
| **Heads in** | 8 | | 1.93 | | 15 | | 7 | 1.93 | | 15 | |
| **Chains in** | 14 | | -0.34 | | 3 | | 14 | -0.34 | | 3 | |
| **Chains out** | 15 | | -0.34 | | 3 | | 13 | -0.32 | | 7 | |
| **Heads out** | 6 | | 1.93 | | 15 | | 7 | 1.83 | | 25 | |

**Supplementary Table 5:** Neutron contrast of the different components used.

| **Compound** | **SLD (10^-6^Å^-2^)** |
| --- | --- |
| dSrf polar group | 2.69 |
| dSrf FA | 0.17 |
| PLPC heads | 1.93 |
| PLPC tails | -0.41 |
| Sitosterol | 0.22 |
| GluCer heads | 2.20 |
| GluCer tails | -0.40 |
