## Supplementary material for "Mechanosensing and Sphingolipid-Docking Mediate Lipopeptide-Induced Immunity in *Arabidopsis*": Material and Methods

**Supplementary Material and methods**

**Purification of surfactin, WLIP and orfamide B**

Surfactin (> 99% purity of a mix of homologues C12/C13/C14/C15 in relative proportions 7/17/45/31%) was purified from spent supernatant of *B. velezensis* liquid culture as previously described^(1)^.

WLIP and orfamide B were produced using *P. putida* COW10 strain and *Pseudomonas* sp., CMR12a*Δsessilin* strain mutant^(2)^, respectively. For purification bacterial cultures were grown in casamino acid media (casamino acid 10 g/L, MgSO_4_ 0.5 g/L, K_2_HPO_4_ 0.3 g/L, pH 7) at 30°C for 48 h on 180 RPM continuous shaking. After 48 h, cultures were centrifuged at 10 000 RPM for 1 h. CLPs were then extracted from cell-free supernatants by acid precipitation. To do so, supernatants were acidified to pH 2 with 4N HCl and incubated at 4°C overnight. The precipitate was collected by centrifugation (14 000 RPM, 1 h), resuspended in water and the pH was adjusted to 8 with NaOH. CLPs were extracted by liquid-liquid extraction (50:50 v/v) (using butanol/ethyl acetate (30:70 v/v) as extraction solvent. Clear solvent phase was further evaporated using rotary evaporator (water bath temperature 55°) and the dried material was resuspended in 100 % ethanol. CLPs were purified by HPLC system (Agilent Series 1100; UV detector 214 nm) by collecting the peaks separated using a C18 column (Luna® Omega 5 μm, 250 x 10 mm) As mobile phase acetonitrile 80% (solution in water) acidified with trifluoroacetic acid (final concentration 0.1%), was used at 5 mL/min (40°C). WILP was purified as a mix of homologues, whereas from orfamide only the major homologue B was purified. Purity of the CLPs was checked by UPLC (Acquity H-class, Waters s.a., Zellik, Belgium) coupled to a single quadrupole mass spectrometer (Waters SQD mass analyzer) on an ACQUITY UPLC® BEH C_18_ 1.7 µm column. Elution was performed at 40°C with a constant flow rate of 0.6 mL/min using a gradient of acetonitrile in water both acidified with 0.1% formic acid as follows: one min at 30%, from 30% to 95% in 3.4 min and maintained at 95 % for 2 min. Compounds were detected in electrospray positive ion mode by setting SQD parameters as follows: cone voltage 120 V, source temperature 130°C; desolvation temperature 400°C, and nitrogen flow: 1000 L.h^-1^.

All CLPs were conserved as powder (-20°C) and freshly prepared as a stock solution (10 mM) in 100 % ethanol on the day of experiments. The 10 µM, concentration used in the experiments (with the exception of Evans blue experiment), was obtained by diluting stock solution in sterile water for plant roots, if not stated otherwise in the methods, and WI_ca_ solution (see protoplasts isolation) for protoplasts treatments. Consequently, the mock treatment consisted in 0.1 % ethanol.

**Plant material and growth condition**

*Arabidopsis thaliana* ecotype Columbia (Col-0) was used as a wild-type control for all plant assays. Seeds were disinfected for 2 minutes in ethanol 75 %, 6 minutes in bleach 5° and rinsed three times in sterilized water. For [ROS]_apo_, [ROS]_intra_ and protoplasts isolation, *Arabidopsis* seeds were grown on half-strength Murashige and Skoog medium (M0222, Duchefa Biochimie; MS) medium with addition of 1% (w/v) sucrose and 14 g/L of agar, for 2 weeks.

For IR experiments and conductivity measurements, one-week-old seedlings, germinated on half-strength MS medium containing 1 % (w/v) sucrose and 14 g/L of agar, were transferred to Araponics systems containing growth solution (0.25% (v/v) FLORAMICRO®, 0.25% (v/v) FLORABLOOM®, 0.25% (v/v) FLORAGRO®; General Hydroponics®) (in volume ratio 1:1:1, as recommended by the manufacturer) where they were grown for four weeks. Growth chamber conditions were constant throughout all experiments, with a photoperiod of 12 hours (100 µmol s^-1^ m^-2^) and a temperature of 23°C.

Mutants of RKs and RLPs *fls2/efr1*, *bak1-5*, *bkk1-1*, *bak1-5/bkk1-1*, *bik1/pbl1*, *cerk1-2*, *sobir1-12*, *sobir1-13*, *dorn1-1*, *lore-5*, *pad3* and *rbohd*, *loh1*, *msl**4/5/6/9/10*, *mca1/2 and* Col-0^aeq^ mutants were described previously^(3–15)^.

**Root Protoplast isolation:**

The protoplast isolation procedure was adapted from^(16, 17)^. Roots from 12 to 15 days-old seedlings were cut into 1-2 mm segments and transferred to protoplasting solution (20 mM MES pH 5.7, 0.4 M mannitol, 20 mM KCl, 10 mM CaCl_2_, 0.1 % (w/v) BSA, 1.5 % (w/v) cellulase R10 (Duchefa Chimie), 0.4 % (w/v) macerozyme R10 (Duchefa Chimie)) for 4 hours at room temperature and in the dark. The suspension was then filtered on gauze to remove root debris and the filtrate was centrifuged for 6 min at 800 RCF. The supernatant was discarded, and the pelleted protoplasts were rinsed once with W5 solution (4 mM MES pH 5.7, 154 mM NaCl, 125 mM CaCl_2_, 5 mM KCl) before being resuspended in WI_Ca_ solution (2 mM MES pH 5.7, 0.5 M mannitol, 20 mM KCl, 2 mM CaCl_2_) at a suitable concentration^(18)^. For experiments with surfactin, protoplasts were used right after their isolation. For experiments with flagellin, protoplasts rested overnight before being used.

**IR evaluation**

Plants grown for 4 weeks in Araponic systems were transferred in 10 mL vials. After 24 hours of rest, plants were transferred to the new vials containing 10 µM Srf or 0.1% ethanol (mock treatment) diluted in the hydroponics solution. After 24 h, plants were inoculated with *Botrytis cinerea* conidia solution. Spores were collected from *B. cinerea* grown on PDA plates for 4 weeks using solution composed of 1,75 g/L KH_2_PO_4_; 0,74 g/L MgSO_4_; 4 g/L glucose and 0.02 % (v/v) Tween 20. After spore collection the concentration was adjusted to 5x10^5^ spores per mL, and spores incubated (30° C, 180 RPM) for 8 hours. Inoculation was conducted by inoculating a drop of 3 µL of conidia solution onto seven leaves per plant. Number of spreading lesions was evaluated 96 h after post inoculation.

**Camalexin quantification**

Camalexin was quantified in plants obtained from IR experiments 96 hours after *B.cinerea* inoculation. Each sample (three samples per treatment) contained five plants (only rosette leaves) pooled together. Plant material was flash-frozen with liquid nitrogen and approximately 100 mg was taken for the extraction. Next, samples were diluted in 1 mL 80 % methanol, agitated (using a bench rotating agitator) at room temperature in the dark for 2 hours and centrifuged (14000 RPM). The supernatant was dried in a rotational vacuum concentrator (2-25 CDplus, Christ) at 50°C, and the pellet was resuspended in 1 mL 100 % methanol, and shaken again for 1 hour. After centrifugation at 14000 RPM, the supernatant was combined with the first one for evaporation. The dry powder was resuspended in 1 mL 100 % methanol.

Samples were then filtered through 0.2 μm PTFE filters before LC-MS based quantification. The analysis was performed using Agilent 1290 Inﬁnity II HPLC system (Agilent) coupled to an accurate mass detector (Jet Stream ESI‐qTOF 6530, Agilent) in positive mode with MS parameters set up as follows : capillary voltage: 3.5 kV; nebulizer pressure: 35 psi; drying gas:8 l min-1; drying gas temperature: 300°C; ﬂow rate of sheath gas: 11 l min-1; sheath gas temperature: 350°C; Nozzle voltage : 1000V; fragmentor voltage: 175 V; skimmer voltage: 65 V; octopole RF: 750 V. Accurate mass spectra was recorded in the range of m/z =100–500. Separation was performed using a C18 Acquity UPLC BEH column (2.1 × 50 mm × 1.7 μm; Waters) and 0.1 % formic acid (solvent A)/acetonitrile acidified with 0.1 % formic acid (solvent B) as mobile phase with constant flow rate at 0.2 mL/min and column temperature set at 40°C. First, solvent B was kept at 25 % during 1 min followed by an increase from 25% B to 60% B in 4 min. Then, 100 % solvent B was applied for 3 min before going back to initial conditions. Masshunter Qualitative Analysis software (Agilent) was used for data analysis. Quantification was performed by comparing camalexin peak area in samples with calibration curve constructed after injection of different concentrations (ranging from 0.08 to 20µM) of pure camalexin standard (Sigma-Aldrich).

**ROS measurements**

Measurement of [ROS]_apo_ in *Arabidopsis* roots was performed by a luminol-based chemiluminescence assay according to^(19)^ and ^(20)^ with some adaptations. Roots were grouped by 10 and separated from the leaves before being cut into small pieces and transferred into a 96-wells white microplate (Lumitrac, Greiner Bio-One, Austria) containing 150 µl of water. The plate was incubated at room temperature in the dark overnight. Then, water was removed and replaced by 90 µl of fresh deionized water before adding 10 µl of a solution containing 200 μM luminol L-012 + 10 μg/ml horseradish peroxydase. Luminescence signal was measured using a Spark® Tecan multiplate reader. First, the background luminescence level was measured every minute during 15min before adding 1µl of Srf 1 mM and flg22 100 µM stock solutions. After the addition of the trigger, luminescence signals were measured every minute for 60 min and results were expressed as % luminescence increase by dividing the measurement at each time point by the measurement at time 0 for each well.

For [ROS]_intra_ measurement, 15 mm long *Arabidopsis* root segments, isolated from different plants, were placed in a well (one root/well) of a microplate (96 Flat Black – Greiner Bio-One™ CellStar™, Fischer Scientific) filled with sterile H_2_O. After overnight incubation, roots were treated with 25 µM DCFH-DA (dichloro-dihydro-fluorescein diacetate; ACROS Organics) for 20 minutes, rinsed with PBS and next, wells were filled with CLP/mock solution. Experiments where [ROS]_intra_ was measured after pre-treatment with the channel blocker or calcium chelator included an additional step where LaCl_3_ (10 mM; Sigma-Aldrich) or EGTA (1mM; Sigma-Aldrich) respectively, were added three minutes before treatments. Fluorescence measurements (excitation wavelength 492 nm, emission wavelength 530 nm) were conducted by a Spark® (Tecan) microplate reader by using nine readings per well. Data expressed as relative fluorescence increase were obtained by subtracting the fluorescence measured at the first time point from the fluorescence measured at each time points (the first time point taken as 0). The fluorescence fold increase was defined for each repeat as the ratio between the fluorescence increase obtained at one time point for treatments and the mean fluorescence increase obtained at the same time point in mock treated tissues.

For [ROS]_intra_ measurements in protoplasts, protoplasts isolated from roots of *Arabidopsis* Col-0 plants were incubated for 10 minutes with 5 µM of DCFH-DA. Then, wells of black 96-well microplates (96 Flat Black - Greiner Bio-One™ CellStar™, Fischer Scientific) were loaded with 150 µL of protoplasts solution per well. After the addition of 50 µL of four times concentrated treatment, the fluorescence was recorded every minute using microplate reader with excitation filter at 485 ± 20 nm and emission filter at 535 ± 25 nm. The data were processed as in roots.

**Calcium influx measurements**

Cytosolic calcium influx ([Ca^2+^]_cyt_) was measured via aequorin-expressing reporter lines (Col-0^AEQ^). For measurements in root tissue, seeds were sterilized with 75% ethanol and grown on 0.5 MS agar plates upright under long day conditions for 16 to 20 days. Roots were cut, pooled in wells of white flat bottom 96-well plates (Lumitrac, Greiner Bio-One, Austria) and incubated in 100 µl of 20 µM coelenterazine-h (p.j.k. GmbH, Germany) for 5 hours in the dark. Luminescence was measured for two minutes before treatment, and 30 minutes after treatment by scanning two rows at a time in 10 seconds intervals using a Luminoskan Ascent 2.1 luminometer (Thermo Fisher Scientific, USA). Each well was discharged for normalization by addition of 150 µL discharge solution (2 M CaCl_2_ in 20 % EtOH) and luminescence was normalized to total luminescence counts remaining (L/Lmax) to obtain [Ca^2+^]cyt.

For measurement in protoplasts, 10 µM of coelenterazine-h (Promega) was added to a protoplast suspension containing 2 x 10^5^ protoplasts. Then, white 96-well microplates were loaded with 100 µL of protoplast suspension and incubated for at least one hour before starting luminescence recording.

The luminescence was recorded by scanning each row in 6 seconds intervals, with an integration time of 300ms, using a Spark® microplate reader (Tecan). The luminescence was first recorded for one minute to obtain resting levels before the manual application of 50 µL of a 3-fold concentrated treatment and a further reading of 25 minutes. To normalize the measurement, the remaining aequorin was discharged by adding 150 µL of 2 M CaCl2 in each well and recording the luminescence for 4 minutes. Data analysis was performed with FlagScreen R-script^(21)^.

Calcium measurements were also performed on protoplasts with the Fluo-4 AM probe. Protoplasts isolated from roots of *Arabidopsis* Col-0, *mca 1/2* or *msl 4/5/6/9/10* mutant were incubated for 1 hour with 5 µM of Fluo-4 AM (ThermoFischer) (from a 5 mM stock solution in DMSO). The suspension was then centrifuged at 750 RCF and the supernatant was discarded to eliminate the remaining free fluo-4 AM. The protoplasts were resuspended in fresh WI_Ca_ solution and were incubated for 1 hour. Microplates (96 Flat Black – Greiner Bio-One™ CellStar™, Fischer Scientific) were loaded with 150 µL of protoplasts solution per well. For experiments with the channel blocker specific to mechanosensitive channels GsMTX-4^(22, 23)^, 7.5 µM of GsMTX-4 was added to protoplasts suspension 10 minutes before the loading in the wells. After the addition of 50 µL of 4 times concentrated treatment, the fluorescence was recorded every 15 seconds using a Spark® microplate reader (Tecan) with an excitation filter at 485±20 nm and an emission filter at 535 ± 25 nm

The values obtained were then converted as normalized fluorescence increase (F/F0) by dividing the fluorescence measured at each time point (F) by the fluorescence measured at the first time point (F_0_).

**Medium alkalinization**

Plants grown as described above were placed in 6 well microplates with roots submerged in the same hydroponic solution used in Araponics systems additionally containing Srf or mock treatment. The change in pH was measured with a pH microprobe (Jenco).

**Conductivity measurement in root medium**

Plants were grown similarly to the ones for the IR experiments, but with prolonged growing time in Araponics (8 weeks). Next, plants were transferred in 10-times diluted MS and rested overnight. For conductivity measurements, 35 mL of root medium was collected and supplemented with 10 µM of Srf, 0.9 % (v/v) Triton X-100 (positive control), or mock treatment. The root of one plant was then immersed in the measurement medium and conductivity was measured using a compact conductivity meter LAQUAtwin-EC-33 (HORIBA scientific).

**Viability test in roots and root protoplasts**

Effect of Srf on root cells was assessed using Evan’s Blue. Roots of six to eight-day-old seedlings were collected and incubated for 30 minutes in a solution containing 0.5 % (v/v) of ethanol (mock treatment), 50 µM of surfactin or 0.9% (v/v) of Triton X-100 (positive control). Next, roots were incubated with 0.25 % (v/v) Evans blue (Sigma-Aldrich) solution for 10 min and rinsed twice with distilled water before microscopic observation.

Effect of surfactin concentration on protoplast viability was assessed with the fluorescent probe fluorescein diacetate (FDA, Sigma Aldrich). Protoplast suspensions were incubated with different concentrations of Srf for 10 min and then incubated with 5µg/mL of FDA (from a stock solution of 5 mg/mL in acetone) for 10 min. Viability of protoplasts was determined with Bürker cell by counting the number of fluorescent protoplasts (viable protoplasts) divided by the total number of protoplasts.

**RNAseq data analysis**

Plants were grown and treated, and RNA was isolated according to^(24)^. Shortly, plant roots were treated for 0, 0.5, 1, 3 and 6 h, each treatment presented as three samples each containing eight roots from different plants. At the end of the treatment samples were flash-frozen in liquid nitrogen and stored at -80°C until the day of RNA extraction. Frozen tissue was homogenised using Eppendorf pestles, and following RNA extraction was conducted using Plant RNeasy Plant Mini Kit (Qiagen, Valencia, CA, USA).

The tool Trimmomatic v0.39^(25)^ was used to trim the raw RNA-seq reads. Quality control on the trimmed reads was performed using FastQC v0.11.8 (Babraham Bioinformatics). We mapped the trimmed reads to the *Arabidopsis thaliana* reference genome (TAIR-10.1) using HISAT2^(26)^. The uniquely mapped reads to the annotated reference genome are estimated as 88.46 %. SAMtools v1.9^(27)^ was applied to generate the required BAM files and their indices. The command line tool featureCounts^(28)^ was employed to calculate the read counts using the latest *Arabidospsis* genome annotation (Araport11_GTF_genes_transposons). Genes with few read counts (<20) were filtered out before further analysis. DESeq2 pipeline (10.1186/s13059-014-0550-8) was used to conduct differential expression analysis with significance parameters set to p<0.05 and log2-fold-change > 2. Genes identification was done using VirtualPlant^(29)^, specificity of DEGs for certain treatments was done using conditional formatting in Excel.

**RT-qPCR experiments**

For RT-qPCR analysis, plants were grown for 12 days on half strength Murashige and Skoog medium (M0222, Duchefa Biochimie) supplemented with 1 % (w/v) sucrose and 1 % (w/v) agar. Seedlings were then transferred individually to 12-wells plates containing Araponics growth solution and grown for 10 additional days in these conditions. Fresh medium was added the day before treatment.

Study of transcriptional change induced by treatments was performed according to^(58)^. Shortly, 22 days-old seedlings were treated with 10 µM surfactin, 100 µg/mL chitin or 0.1 % ethanol (negative control) for 6 h. Each treatment consisted of three samples each containing eight roots. At the end of the treatment, root tissues were flash-frozen in liquid nitrogen and homogenized using Eppendorf pestles. RNA extraction was performed using Plant RNeasy Plant Mini Kit (Qiagen, Valencia, CA, USA). RT-qPCR reactions were conducted using the Luna® Universal One-Step RT-qPCR Kit (New England Biolabs, Ipswich, MA, United States) following the manufacturer’s instructions. The thermal cycling program applied on the ABI StepOne was: 55 °C for 10 min, 95 °C for 1 min, 40 cycles of 95 °C for 10 s and 60 °C for 1 min, followed by a melting curve analysis performed using the default program of the ABI StepOne qPCR machine (Applied Biosystems). The real-time PCR amplification was run on the ABI step-one qPCR instrument (Applied Biosystems) with software version 2.3. Primers used are listed in **Supplementary Table 2**. The relative gene expression analysis was conducted by using the 2^-ΔΔCt^ method with the *UBQ5* gene^(59)^ as a housekeeping gene to normalize mRNA levels between different samples.

**Liposome preparation**

1-palmitoyl-2-linoleoyl-sn-glycero-3-phosphocholine (PLPC), β-sitosterol (Sito) and D-glucosyl-ß-1,1'-N-palmitoyl-D-erythro-sphingosine (GluCer) were purchased from Avanti Polar Lipids and used without further purification. Large unilamellar vesicles (LUVs) were prepared for ITC, AFM, WAXS and Laurdan generalized polarization experiments. The different lipid mixtures (PLPC (for ITC), PLPC-Sito (80-20 molar ratio) (for ITC), PLPC-GluCer (80-20 molar ratio) (for ITC) and PLPC-Sito-GluCer (60-20-20 molar ratio) (for ITC, AFM, WAXS and Laurdan generalized polarization)) were dried from a chloroform/methanol (Scharlau Lab Co.) (2/1, v/v) solution under reduced pressure in a rotary evaporator at 30°C and then kept under vacuum overnight. The dried lipid films were then hydrated in Tris 10 mM NaCl 150 mM buffer at pH 8.5 (to 5 mM of lipid for ITC, 2 mM for AFM and to the final 4% total concentration for WAXS) or in MES 10 mM NaCl 150 mM buffer at pH 5.8 (to 1 mM of lipid for Laurdan analysis) during 1 h at 45°C with vortex mixing applied every 15 min and then subjected to five freeze/thaw cycles.

The dispersions were finally extruded fifteen times through two stacked Nuclepore 100 nm polycarbonate filters using a Lipex Biomembranes (Vancouver, BC) extruder to obtain LUVs (112.2 ± 3.6 nm). The average size of LUVs was determined at 25°C by dynamic light scattering (DLS) method using a Zetasizer nano ZS (Malvern instruments, UK) with a He–Ne laser source at a wavelength of 633 nm. The scattered light intensity was measured at a scattering angle of 173°.

For FT-IR experiments, multilamellar lipid vesicles (MLVs) were prepared as described^(30)^. PLPC, Sito and GluCer (60-20-20, molar ratio) were dissolved in a chloroform /methanol mixture (2/1, v/v) alone or in the presence of Srf at a lipid-to-Srf molar ratio of 90-10. The chloroform/methanol mixture was evaporated to obtain a lipid film dried under vacuum overnight. The resulting film was hydrated by deuterium oxide above the phase transition temperature of lipids. The hydrated film was vortexed continuously to obtain MLVs.

**ITC analysis**

ITC analyses were performed with a VP-ITC Microcalorimeter (Microcal, Northampton, USA). The calorimeter cell (Volume of 1.4565 mL) was filled with a 10 µM (below the CMC concentration) Srf solution in buffer (Tris 10 mM, NaCl 150 mM at pH 8.5). The syringe was filled with a suspension of LUVs at a lipid concentration of 5 mM. A series of 10 µL injections was performed at constant time intervals (6 min) at 25°C. The solution in the titration cell was stirred at 305 RPM. Prior to each analysis, all solutions were degassed using a sonicator bath. The heats of dilution of vesicles were determined by injecting vesicles in buffer and subtracted from the heats determined in the experiments. Data were processed by software Origin 7 (Originlab, Northampton, USA). All measurements were repeated at least three times with two different vesicle preparations.

**Hypermatrix calculation**

The Hypermatrix method^(31, 32)^ is a simple docking method that allows for the calculation of the interaction between two molecules. The molecule of interest (Srf in this study) was fixed at the center of the system and oriented at a hydrophobic (pho)/hydrophilic (phi) interface using the TAMMO procedure^(32)^. The lipid molecule was also oriented at the pho/phi interface and was positioned around the central Srf by rotations and translations (more than 10 million positions were tested). For each position, an energy value was calculated, according to a home-designed force field^(33)^. The energy values together with the coordinates of all assemblies were stored in a matrix and classified, according to decreasing values. The first stable match was considered as the best assembly between the two molecules.

**Molecular dynamics simulation**

Surfactin SC14 was studied by molecular dynamics (MD) in presence of a membrane of PLPC and PLPC-Sito-GluCer (60-20-20) with the Gromacs v4.5.4 software^(34)^. Coarse-grained simulations have been carried out first for Srf insertion and building of the lipid membrane. Models were converted to a CG representation suitable for the MARTINI 2.1 forcefield^(35)^ with the Martinize script and a coarse-grained Srf was placed over the membranes with the insane tool^(36)^. This first insertion was used to place afterwards 9 Srf molecules in both leaflets at the same membrane level. Water particles were then added as well as ions to neutralize the system. A 2000-steps steepest-descent energy minimization was performed to remove any steric clashes. An equilibration of 100 ns with a 20 fs time step has been carried on. Temperature and pressure were coupled at 300 K and 1 bar using the weak coupling Berendsen algorithm^(37)^ with τT = 1 ps and τP = 1 ps. Pressure was coupled semi-isotropically in XY and Z. Non-bonded interactions were computed up to 1.2 nm with the shift method. Electrostatics were treated with ε = 15. The compressibility was 10^-5^ (1/bars). The system was then transformed to an atomistic resolution with backwards^(38)^. Atomistic simulations have been performed with the GROMOS96 54a7 force field^(39–41)^. Parameters of the ester bond between the acyl and LEU7 residues and for the acyl chain were taken from the GROMOS96 54a7 force field^(42)^. Parameters for the GluCer was taken from^(43)^. All the systems studied were first minimized by steepest descent for 2000 steps. Then NVT and NPT equilibrations were carried on for 0.1 and 1 ns. Srf was under position restraints and periodic boundary conditions (PBC) were used with a 2 fs time step. Production runs were performed for 1µs. All the systems were solvated with SPC water and the dynamics were carried out in the NPT conditions (300 K and 1 bar). Temperature was maintained by using the Nose-Hoover method (44) with τT = 0.5 ps and a semiisotropic pressure was maintained by using the Parrinello-Rahman barostat^(45)^ with a compressibility of 4.6 × 10^-5^ (1/bar) and τP = 5 ps. Nonbonded interactions were evaluated using a twin-range cutoff scheme. Interactions within the shorter-range cutoff (0.8 nm) were calculated every step, whereas interactions within the longer cutoff (1.4 nm) were updated every 5 steps, together with the pair list. In all the simulations, a reaction-ﬁeld correction was applied to the electrostatic interactions beyond the long-range cutoff^(46)^ using a relative dielectric permittivity constant εRF of 62. Bond lengths were maintained with the LINCS algorithm^(47)^. The trajectories were performed and analyzed with the GROMACS 4.5.4 tools as well as with homemade scripts and softwares, and 3D structures were analyzed with both PYMOL (DeLano Scientific, http://www.PyMOL.org) and VMD softwares^(48)^.

**Neutron Reflectivity (NR)**

For deuterated Srf production, *Bacillus velezensis* GA1 was cultured in 20 mL Liquid LB broth supplemented with 1g/L of deuterated Leucine (Cambridge isotope) for 24 h at 30°C under agitation. Extraction was performed by liquid-liquid partitioning using 20 mL of ethyl acetate/butanol (70:30 v/v). Solvents were evaporated under rotary evaporation and powder was resuspended in 2 mL of 100% ethanol prior to UPLC (Agilent 1290 Inﬁnity II) purification by collecting each deuterated C15 Srf peak. 10 µL extract was separated using C18 column (C18 Acquity UPLC BEH column; 2.1 × 50 mm × 1.7 μm; Waters) An isocratic 0.2 ml/min flow with 75 % acetonitrile/water (acidified with 0.1 % formic acid) was applied for 10 min before raising up to 100 % acetonitrile for 5 min and going back to initial ratio before the next injection. Purified deuterated Srf structure was confirmed by LC-MS/MS using the same LC method coupled with an accurate mass detector (Jet Stream ESI‐Q‐TOF 6530) in positive mode with parameter set up as follows: parameters: capillary voltage: 3.5 kV; nebulizer pressure: 35 psi; drying gas:8L/min; drying gastemperature:300°C; ﬂow rate of sheath gas: 11 L/min; sheath gas temperature: 350°C; fragmentor voltage: 175 V; skimmer voltage: 65 V; octopole RF: 750 V. Collision energy 40V. Quantification was performed by comparing peak area of purified compound with the peak area of a commercial standard (Lipofabrik, Villeneuve d’Ascq). The relative amounts of the deuterated surfactin C15 is presented in **Suppl. Table 3**.

Lipid bilayer (ternary mixture of PLPC-Sito-GluCer – 60-20-20 molar ratio) depositions on silicon for NR measurements were obtained by injecting directly in the measuring cell (6mL) at room temperature the extruded vesicles prepared as for X-ray investigation but to the final concentration of 0.5mg/ml, according to the procedures described in^(49)^.

Neutron reflectivity data were acquired at the MARIA neutron reflectometer^(50)^ operated by Jülich Centre for Neutron Science at Heinz Maier-Leibnitz Zentrum in Garching (Germany), using custom temperature-regulated liquid cells^(51)^. The measurements were performed using two different wavelengths, 10 Å for the low-q region and 5 Å for the high-q region up to 0.25 Å^-1^, with a 10% wavelength spread. The change of solvent contrast in the liquid cells was performed using a combination of valves and a peristaltic pump, at small flow rates ~ 0.5 ml/min. In a reflectivity experiment a grazing beam is sent to the sample and the reflected intensity is collected as a function of the reflection angle momentum transfer perpendicular to the interface qz (qz = 4 λ sin ϑ /2, where ϑ and λ are the angle of the incident beam and wavelength, respectively). The technique allows to get information about the sample cross structuring in a non-invasive way^(52, 53)^. The silicon oxide layer, the water layer between the silicon oxide and the membrane and the different hydrophilic and hydrophobic layers of the lipid membranes have been modelled as defined layers with a proper thickness, compactness and mean composition (and therefore contrast to neutrons). Reflectivity has been measured from the silicon supports and the samples in different water solutions (H_2_O and D_2_O - Sigma Chemical Co). After bare membrane characterization in two solvents, the Tris 10 mM NaCl 150 mM buffer at pH 8.5 solution was injected into the cell and NR measure was performed. Finally, 2 µg of Srf in Tris HCl buffer at pH 8.5 have been injected into the cell, to the final 95:5 membrane:Srf molar proportion. NR measurements have been performed on this system after 30 minutes incubation. Data have been analyzed by the MotoFit program^(54)^. Data relative to the same system measured in different water contrasts have been analyzed by contemporary fits.

The scattering length density (SLD) of the mixture used has been calculated according to relative molar proportions, by considering a volume of 1143 Å^3^ for the hydrophilic portion and of 373 Å^3^ for the fatty acid (FA) ^(55, 56)^ (**Suppl. Table 4 and 5**).

**Wide Angle X-Ray Scattering (WAXS)**

LUVs (PLPC-Sito-GluCer (60-20-20 molar ratio) were separated on two aliquots: to the first, the required volume of solution of Srf in buffer was added to obtain a final lipid-to-Srf 95-5 molar proportion, while the second was diluted with the same amount of pure buffer.

WAXS investigations were performed at ID02 beamline of the European Synchrotron (ESRF, Grenoble, France). The sample-to-detector distance for SAXS was 1 m to cover a q-range, 0.07<q <5 nm^-1^, being q the scattering wave vector defined as q = (4π/λ) sin ϑ/2, with λ the wavelength (λ ∼ 1 Å) and ϑ the scattering angle. Measurements were performed using polycarbonate capillaries of 2 mm thickness (ENKI, Concesio, Italy) as sample containers. The measured patterns were corrected for detector artefacts, normalized to absolute intensity scale and azimuthally averaged to obtain the intensity profile I(q) using standard procedures^(57)^. For each sample, 3−4 frames were acquired which were subsequently averaged after excluding any possible radiation damage. The background scattering of the buffer was also measured. The averaged background signal was subtracted from each averaged sample intensity profile and best fits for the measured data were derived using the model of spherical core-shell objects as implemented in the SAXSutilities analysis package [http://www.sztucki.de/SAXSutilities/].

**Infrared spectroscopy (FTIR)**

Infrared spectra were measured with 128 scans at 4 cm^−1^ resolution using a Bruker Equinox 55 spectrometer (Karlsruhe, Germany) equipped with a liquid nitrogen-cooled DTGS detector and linked to a computer with OPUS software marketed by Bruker. During all measurements, the spectrometer was continuously purged with N_2_ flux. All the experiments were performed with a demountable cell (Bruker) equipped with CaF_2_ windows on which MLVs were deposited. Each spectrum is the representative of at least two independent measurements.

**Atomic force microscopy experiments**

Supported lipid bilayers (SLB, ternary mixture of PLPC, Sito, GluCer – 60-20-20 molar ratio) were reconstructed on freshly cleaved mica substrates by allowing the fusion of a 2 mM LUVs solution (V = 100 µL) at 55°C for 45 min. Samples were then let for thermalization at ambient temperature for 30 min without dewetting and immersed in 3 mL of 10 mM Tris 150 mM NaCl buffer at pH 8.5. Not to damage the samples, AFM images were obtained in the quantitative imaging (QI) mode of a JPK Nanowizard III setup, with a minimal applied force of 200 pN and a speed of 50 µm/s. Soft sharpened silicon nitride cantilevers (MSCT, Bruker) were used and calibrated before any experiment using the thermal noise method (k ~ 0.02 N/m). Srf, prepared in 10 mM Tris 150 mM NaCl buffer at pH 8.5, was finally injected at a final concentration of 3 µM. AFM Images were then recorded at different time points in different areas to follow its impact on the lipid bilayer.

***Laurdan polarization on root protoplasts and lipid vesicles***

Protoplast suspension (1-2 10^5^ protoplasts / mL) or 100 µM LUVs preparation was incubated with 2 µM of Laurdan (Sigma-Aldrich) for 1h. Then, wells of black 96-well microplates (Greiner Bio-One™ CellStar™, Fischer Scientific) were loaded with 100 µL of protoplasts or LUVs solution per well. The fluorescence was recorded using Spark® microplate reader (Tecan) by performing fluorescence scan between 405 and 520 nm with an excitation wavelength at 360 nm. The fluorescence was recorded once before treatment and at various times after the addition of 25 µL (for protoplasts) or 100 µL (for LUVs) of treatment. The treatments were prepared in the buffer of protoplasts (WI_Ca_ solution) or LUV buffer respectively with a concentration taking into account the dilution factor occurring at the addition of treatment into 100 µL of protoplasts or LUVs solution (5-times more concentrated for protoplasts and 2-times more concentrated for LUVs). The generalized polarization (GP) was defined as GP = $\frac{{(I}_{440nm}-I_{490nm})}{{(I}_{440nm}+I_{490nm})\text{ }}$, were I_440nm_ and I_490nm_ represents the blank-subtracted fluorescence intensities at emission wavelengths of 440 nm and 490 nm respectively. Variation of GP (ΔGP) is defined as the subtraction of GP measured at each time point following treatment and GP measured before treatment.

**Statistical methods**

Statistical details of experiments are specified in the figure legends. All statistical analyses were performed in GraphPad Prism 8.0.1.
